# A tissue-scale strategy for sensing threats in barrier organs

**DOI:** 10.1101/2025.03.19.644134

**Authors:** Diep H. Nguyen, Jiakun Tian, Sean-Luc Shanahan, Connie Kangni Wang, Tyler Jacks, Xiao Wang, Pulin Li

## Abstract

Barrier organs rely on a limited set of pattern recognition receptors (PRRs) to detect diverse immunogenic challenges. How organs assess threats and adjust immune responses to balance host protection with collateral tissue damage remains unclear. Here, by analyzing influenza infection in the lung using single-molecule imaging and spatial transcriptomics, we discovered a tiered threat-sensing strategy at the tissue scale, where the probability of detecting and responding to infection is lowest in the outermost epithelia and highest in the inner stroma. This strategy emerges from spatially graded PRR expression that results in cell-type-specific probabilities of threat-sensing across the tissue, a design broadly adopted by barrier organs. Selectively increasing PRR expression in lung epithelia *in vivo* exacerbated tissue damage upon inflammatory challenge. These results reveal a spatially tiered strategy to tolerate threats restricted within the epithelia, and yet enable progressively potent immune responses as threats invade deeper into the tissue.

## Introduction

When facing diverse environmental insults, organs need to carefully balance the objectives of eliminating pathogens by activating the immune system versus maintaining tissue integrity by limiting the magnitude and duration of such activation. Premature, insufficient, and excessive activation of immunity can all cause severe pathology ^1–3^. How barrier organs, which directly interface the environment, execute this delicate balance at each step of immune activation to optimize their fitness remains a mystery. Understanding this question is particularly important for the lung, an organ that is vulnerable to both pathogens and tissue damages, yet remarkably quiescent compared to other barrier organs ^4,5^.

Detecting pathogens through pattern-recognition receptors (PRRs) is the initial step that triggers the complex immune cascade in response to pathogen infection. PRR activation induces the secretion of cytokines and chemokines that mobilize the immune system, making PRR activation an appealing knob for regulating the timing and level of immune activation ^6–8^. As the barrier cell type in the lung, epithelial cells are often the initial targets of infection and therefore the central focus for studying PRR pathway activation ^9–11^. However, individual infected lung epithelial cells lack the capability to scale the amount of cytokines produced based on the level of threat, such as the viral load. In fact, the production of type I interferons (IFN), a major consequence of PRR pathway activation, is highly stochastic, raising the question of how cells in the lung collectively tune the level of cytokines during initial infection ^12–20^.

Furthermore, highly virulent and opportunistic pathogens can often escape detection by epithelial cells, breach the barrier, and invade the stromal compartment to infect cells, such as fibroblasts, tissue-resident immune cells, and endothelial cells ^21,22^. Curiously, PRR pathways appear to be ubiquitously present in most cell types, suggesting that all cells have the potential to sense pathogens ^8^. However, the contribution of cells in the stromal compartment to the initial events of pathogen detection and immune activation has not been systematically analyzed and compared to the epithelial layer. These observations raise the central question of how the pathogen-sensing and immune-activation function is spatially organized within the lung.

By quantitatively analyzing the early phase of influenza A virus infection in the mouse lung at single-cell resolution, we discovered a spatially tiered threat-sensing strategy that helps balance host protection with collateral tissue damage. Specifically, different cell types differed drastically in the probability of detecting the virus and activating PRR pathways, although they were all equipped with PRRs. Counterintuitively, the outermost airway and alveolar epithelial cells had the lowest probability, and the innermost endothelial cells had the highest probability, primarily due to variations in the expression level of PRRs. When this spatially tiered design was perturbed by epithelia-specific overexpression of retinoic acid inducible gene I (RIG-I), the tissue suffered from prolonged inflammation, more severe damage, and delayed regeneration upon pathogen-free inflammatory challenge. Together, we discovered a tissue-scale strategy that operates at the initial step of threat sensing and immune activation. It allows tissues to assess the degree of threat and adapt the immune responses, from tolerating minor epithelia-restricted threats to efficiently activating self-defense against severe infection that invades the tissue. This appears to be a universal principle across nucleic-acid sensing PRR pathways and across barrier organs.

## Results

### Cells in the lung detect infection in a probabilistic manner

To understand how the lung assesses the level of threat and mount the appropriate responses at the initial step of immune activation, we used the mouse model of influenza A WSN H1N1 virus infection, which infects both airways and alveoli in mice, the two major anatomical structures in the lung ^23,24^. Spatially restricted infection foci formed in both airways and alveoli within 36 hours post infection (hpi) (**Fig. 1A, S1A, B**). Using single-molecule hybridization chain reaction RNA fluorescence *in situ* hybridization (smHCR) to quantify influenza *NP* (a proxy for viral load) and *Ifnb1* (a proxy for IFN produced by infected cells upon viral infection ^25^) in single cells, we found that only ∼5% of all infected cells expressed detectable levels of *Ifnb1* transcripts (**Fig. 1B**). Furthermore, the level of IFN per cell was not regulated by the viral load; instead, the viral load controlled the probability of producing IFN in an infected cell (**Fig. 1B, C**). A similar probabilistic control of IFN production was further confirmed in human lung epithelial A549 cells, regardless of the specific *IFN* transcripts or viral genetic components measured (**Fig. S1C-E**). To eliminate the paracrine effect of IFN on how neighboring cells respond to infection ^26^, we blocked JAK/STAT signaling downstream of IFN with Ruxolitinib and found the probabilistic control model held true (**Fig. S1F-H**).

**Figure 1.**
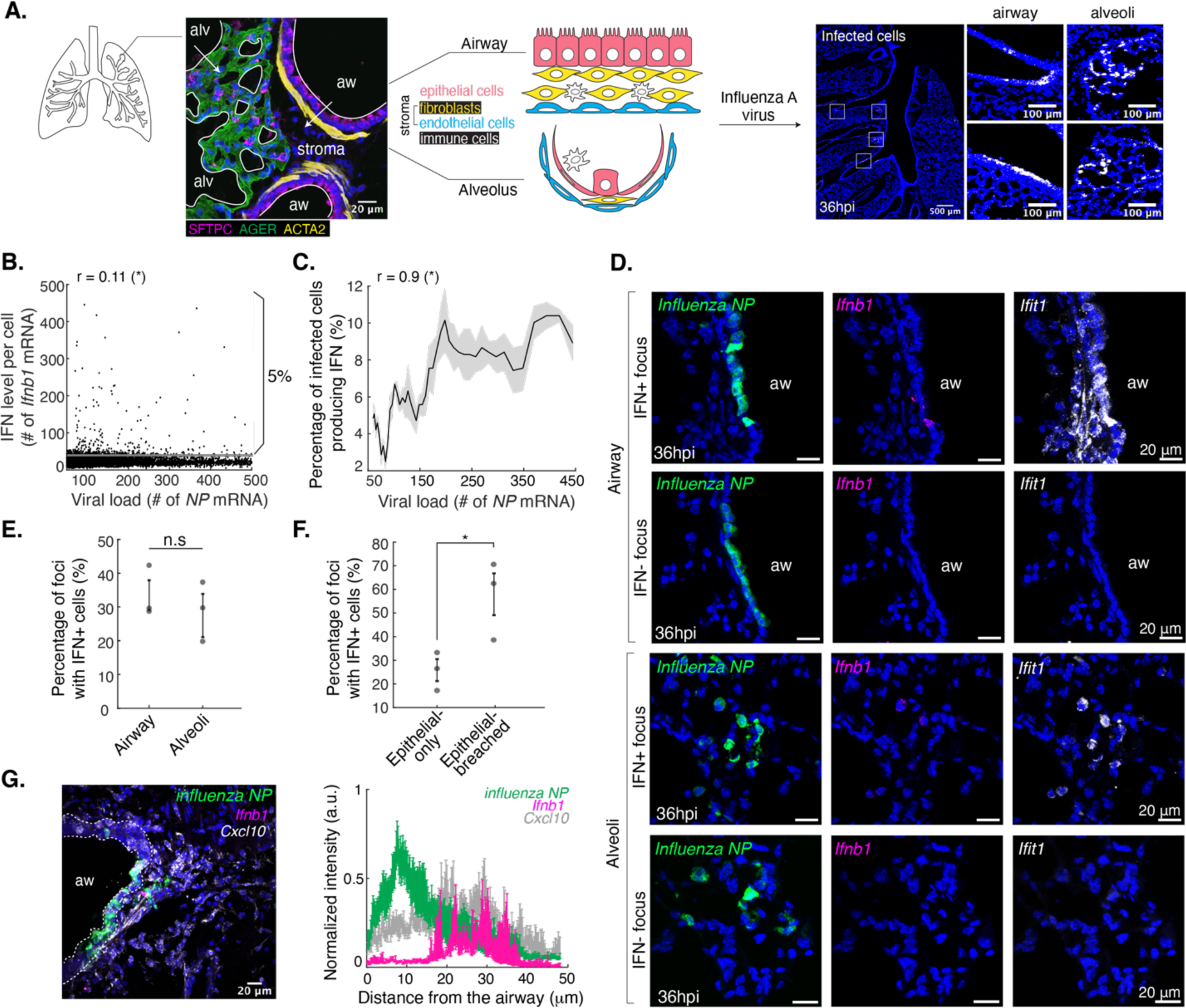
Cells in the lung detect infection in a probabilistic manner. **A.** Representative images and schematic of two distinct barrier structures in murine adult lung labeled with antibodies against SFTPC (alveolar type 2 and distal airway epithelia), AGER (alveolar type 1), ACTA2 (smooth muscle). The outermost layer of the lung tissue (*solid white line*) is lined by airway and alveolar epithelial cells. The stromal compartment underneath comprises fibroblasts, endothelial cells, and tissue-resident immune cells. Influenza A virus WSN H1N1 infects both barrier structures at 36 hour post infection (hpi). **B.** Lack of correlation between mouse *Ifnb1* mRNA and viral load at the single-cell level in infected lungs. Threshold (*solid gray line*) was determined based on the number of transcripts counted in uninfected lungs. **C.** The percentage of IFN+ cells positively correlated with viral load in infected lungs. Infected cells were sorted and binned based on viral load, and the percentage of IFN+ cells was calculated for each viral load bin. Shaded region indicates mean +/- SEM. For **B** & **C**, n = 10651 cells pooled from 3 mice, and r indicates Spearman’s rank correlation coefficient. Asterisk denotes statistical significance. **D.** Representative single-molecule HCR-RNA-FISH (smHCR) images of influenza-infected foci in airway and alveoli at 36 hpi. A focus was chosen as a cluster of infected cells within a 0.15mm x 0.15mm field of view. aw, airway. **E.** Similar percentage of foci with IFN+ cells in the airway and alveoli. For **E** & **F**, each dot represents an animal and error bars indicate mean +/- SEM. Asterisk denotes statistical significance, and n.s. denotes no significance using one-way ANOVA test. **F.** The percentage of foci with IFN+ cells was higher in epithelial-breached foci than epithelial-only foci. **G.** Quantifying the spatial distribution of IFN+ cells along the exterior-interior axis of the airway. A representative smHCR image of an epithelial-breached focus in the airway (*top*). Dotted lines mark the epithelial border. Normalized intensity of influenza and host mRNA as a function of distance from the airway (*bottom)*. n = 10 foci.

The low probability of infected cells producing IFN led to drastic heterogeneity among foci, with only 40% of the foci being IFN+ with 1-2 cells producing IFN at 36 hpi (**Fig. 1D, E**). By simultaneously probing for interferon-stimulated genes (ISG), such as *Ifit1* and *Isg15,* to identify foci that have experienced IFN, we confirmed the high detection efficiency of *Ifnb1* transcripts, with >90% of ISG+ foci having at least one IFN+ cells (**Fig. S1I**) ^27^. Together, these observations demonstrate that infection within 36 hpi does not guarantee immune activation at the single-cell level, with a higher probability of IFN production at a higher viral load. This suggests that an infected tissue can tune the degree of immune response to infection severity by modulating the fraction of infected cells producing IFN.

### Cell types in the stromal compartment are more likely to detect infection and produce IFN than epithelial cells

Because influenza first attacks epithelial cells, we asked if the IFN+ vs IFN-infection foci mainly differed in the response of epithelial cells to infection. We focused on the airway foci due to the distinct separation between epithelial and stromal tissues. We found that larger foci exhibited viral breaching of the epithelial barrier and infection of cells within the stromal compartment, and these breached foci were also more likely to be IFN+, compared to epithelia-only foci (**Fig. 1F, S1J**). Furthermore, we found that IFN+ cells were localized in the stromal compartment, even though most of the infected cells were airway epithelia, suggesting that non-epithelial cell types might be more effective at detecting and responding to infection than epithelial cells (**Fig. 1G**).

To comprehensively characterize the cell types infected in the lung and quantify the probability of IFN production for each infected cell type, we profiled influenza-infected mouse lung using STARmap PLUS ^28^ (**Fig. 2A**). Using influenza NP protein to identify infection foci, we sequenced 10 lung sections, which included 89 infection foci, and their surrounding regions from different anatomical locations (30 airway foci and 59 alveolar foci). A curated list of 237 genes were used to probe markers for cell types, infection status, and immune response status (**Supplementary Table 1**). We identified most types of epithelial, fibroblast, endothelial, and immune cells according to the previously reported markers (**Fig. 2B**, **S2A-C**).

**Figure 2.**
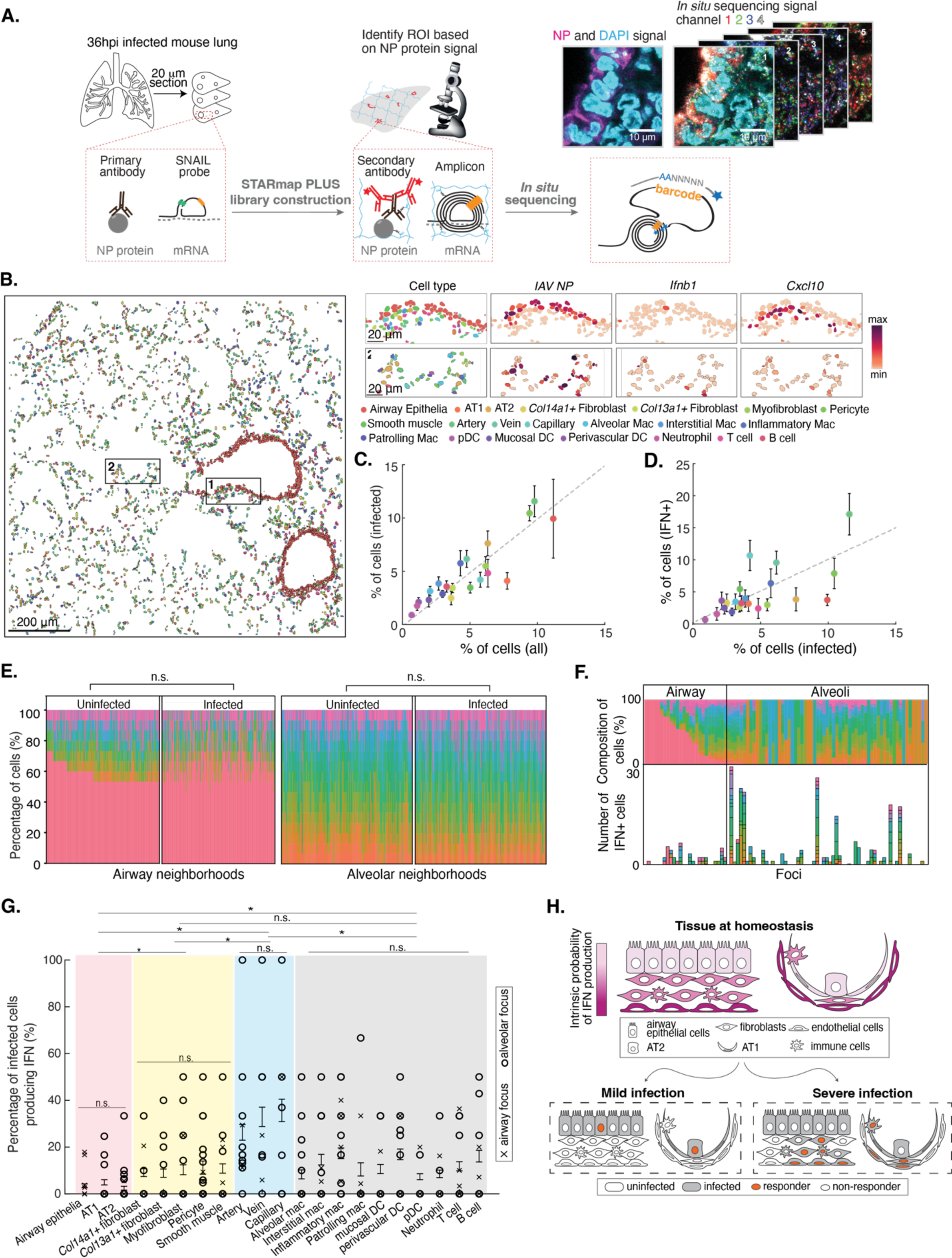
Cell types in the stromal compartment are more likely to detect infection and produce IFN than epithelial cells. **A.** Schematics of STARmap PLUS on 20 μm influenza-infected mouse lung sections at 36 hpi. Representative images show the simultaneous mapping of cell nuclei via DAPI, NP protein signal, and cDNA amplicons. Five cycles of *in situ* sequencing were performed to decode the amplicon barcodes. ROI, region of interest. **B.** STARmap identified cell types, infection state, and signaling states in influenza-infected lungs. Insets: zoom-in on boxed regions with examples of cell types, infection, and immune response status. Each polygon represents a cell, color-coded by cell type. AT1, alveolar type 1; AT2, alveolar type 2; mac, macrophage; DC, dendritic cell; pDC, plasmacytoid dendritic cell. **C.** Most cell types had similar susceptibility to influenza infection. Dotted line represents the expected likelihood of being infected given the relative abundance of the cell type in the tissue. For **C**, **D**, **E**, **F**, colors correspond to cell types in **B**. Each dot represents a cell type. Error bars indicate mean +/- SEM across 10 tissue sections. **D.** Endothelial cell types were over-represented and epithelial cells were under-represented among IFN+ cells. IFN+ cells were defined by the combined expression of *Ifnb1, Ifna4,* and *Ifnl2* transcripts. Dotted line represents the expected likelihood of producing IFN given the relative abundance of the cell type among infected cells. **E.** Composition of cell types in airway and alveolar neighborhoods did not change after infection at 36 hpi. Each column is a neighborhood. n.s. denotes no significance using Fisher exact test. **F.** Most cell types contributed to the production of IFN stochastically. Cell-type composition of infected cells (*top*) and that of IFN+ cells (*bottom*) in each focus. Each column is a focus. **G.** The percentage of IFN+ cells among infected cells by cell type. Each point represents a measurement per focus. Error bars indicate mean +/- SEM across foci. Asterisks denote statistical significance, and n.s. denotes no significance using Kruskal-Wallis test. **H.** Tiered threat-sensing model: Spatially graded probability of IFN production upon infection along the exterior-interior axis of the organ, which allows the tissue to adapt the level of immune responses to varying severity of infection.

We reasoned that the contribution of different cell types to the overall level of IFN depends on the relative abundance of the cell type, its susceptibility to influenza infection, and its intrinsic probability to produce IFN. By comparing cell-type composition between all the cells and infected cells pooled from all sections, we observed similar infection susceptibility among different cell types, consistent with a previous observation using single-cell RNA-seq ^29^ (**Fig. 2C**). Of these infected cells, structural cell types, namely epithelial, fibroblast, and endothelial cells, made up the majority of IFN-producing cells (**Fig. 2D**). Notably, endothelial cell types (artery, vein, and capillary endothelia) were over-represented among IFN-producing cells, whereas epithelial cell types (airway epithelia, AT1, and AT2) were under-represented among IFN-producing cells. The results revealed that different cell types in the lung have similar susceptibility to influenza A virus infection, but different probability of producing IFN upon infection.

To understand how infection and IFN production played out spatially in heterogeneous tissues, we analyzed cell types and states in individual focus. We first examined the cell-type composition among neighborhoods, defined as a collection of 14 nearest neighbors around a center cell, and found no significant change in cell-type composition upon infection (**Fig. 2E**, **S2D**). Structural cells made up 80% of most neighborhoods and no major immune infiltration occurred at 36 hpi with the viral dose we used. We then analyzed foci, defined as spatially restricted clusters of infected cells, and found that 55 foci (61%) were IFN+/ISG+ in both airway and alveoli (**Fig. 2F, S2E**). Larger foci with breached epithelia had a higher probability of being IFN+, consistent with our previous smHCR results (**Fig. S2F-G**). Within these IFN+ foci, although most cell types can contribute to IFN production, endothelial cell types made up the majority of IFN-producing cells (**Fig. 2F**). Furthermore, we found that endothelial cell types had the highest probability of producing IFN (30-40%) and epithelial cell types had the lowest probability (∼5%), with fibroblast and immune cells in between (∼15%) (**Fig. 2G**). Notably, cell types belonging to the same family (e.g. airway and alveolar epithelial cell types within the epithelial cell family) did not differ significantly in their probability (**Fig. 2G**). These results suggest that tissue structural cells are most likely IFN producers in the early phase of influenza infection, and different families of structural cells produce IFN at drastically different probability.

Together, our data reveal an unexpected spatial organization of an organ’s innate immunity: the probability of IFN production is spatially graded, with the lowest probability in the outermost epithelial layer, including both airway and alveolar epithelia, and highest probability in the innermost endothelial layer (**Fig. 2H**). Instead of tuning the amount of IFN produced per cell, the tissue modulates the magnitude of immune response by tuning the probability of producing IFN (**Fig. S2H**). This design enables a tissue-scale strategy to deal with threats of varying degrees. When the infection is restricted to the epithelia, the tissue limits the number of cells that produce cytokines to curb the immune response. But when the infection gets more severe and invades the deeper tissue, infected cells are more likely to detect infection and produce cytokines to mobilize the immune system.

### Differential probabilities of detecting infection are intrinsic properties of cell types

The probability of producing IFN by an infected cell could depend on both the virus and the state of the host cell. To ask if the differential probability of producing IFN among different cell types was due to different viral loads, we quantified viral *NP* mRNA level using the STARmap data and found similar viral loads among different cell types, with the exception of airway epithelial cells, alveolar type II cells, and alveolar macrophages (**Fig. S2I**). Although these three cell types harbored higher viral load, they had lower probability of producing IFN, further suggesting the intrinsic properties of host cells play a more significant role in determining the probability of IFN production.

To better quantify the intrinsic capability of viral detection and cytokine production among different cell types, we performed synchronized viral infection in cultured primary human lung epithelial, fibroblast, and endothelial cells *ex vivo*. The cell culture system allows us to circumvent several confounding factors associated with infection in the lung, including that the different cell types are infected at different time points and consequently, previously infected cells might influence how other cells respond to infection. In particular, IFN paracrine signaling can induce a positive feedback that enhances the antiviral state of the cells, and such effect can be blocked by a JAK/STAT signaling inhibitor Ruxolitinib ^26^ (**Fig. S1F**). By applying the same viral dose and Ruxolitinib to all cell types, we observed that epithelial cells had the lowest probability of producing IFN (∼8%), and endothelial cells had the highest probability (30%), with fibroblasts in between the two (20%), consistent with our *in vivo* results (**Fig. 3A-C**). Surprisingly, this trend persisted even among human epithelial, fibroblast and endothelial cells from different organs, such as immortalized lung epithelial, dermal fibroblast, and umbilical vein endothelial cells (**Fig. 3D, E**). Even though the range of viral loads varied slightly across cell types, at the same viral load, different cell types produced IFN at different probability (**Fig. 3F**). Together, these results show that the probability of IFN production is an intrinsic feature of each cell type, and cell types belonging to each of the broader cell families, namely epithelial, fibroblast, and endothelial cells, have similar likelihood of producing IFN (**Fig. 2G**). This conserved feature across cell types suggests the existence of generalizable mechanisms that regulate the probability of IFN production.

**Figure 3.**
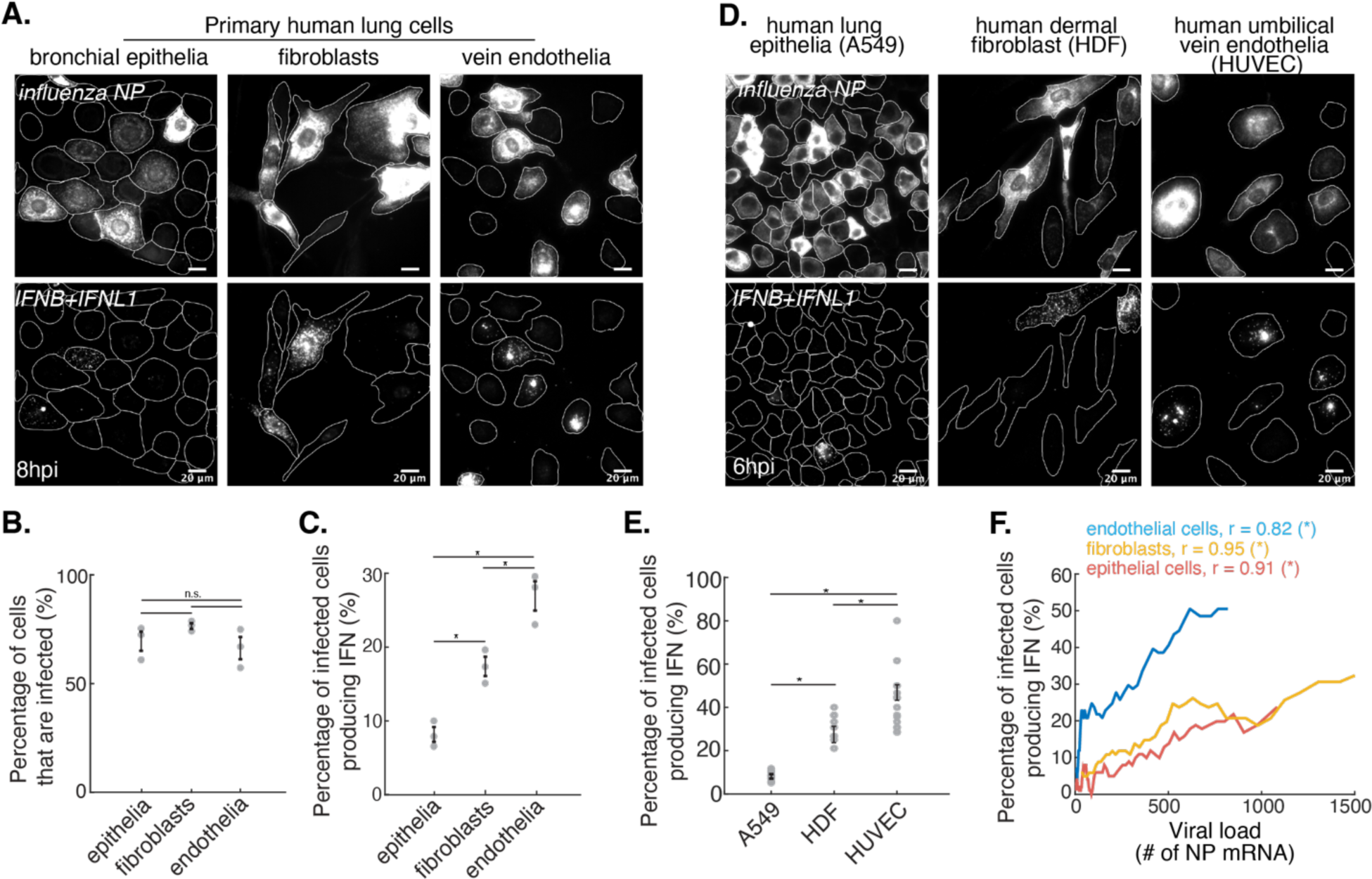
Differential probabilities of detecting infection are intrinsic properties of cell types. **A.** Representative smHCR images of influenza-infected human primary lung cells at 8 hpi. **B.** Infection efficiencies were comparable across primary human lung cell types. Each dot represents an experimental replicate. Average n = 2988 (epithelial cells), 733 (fibroblasts), 1314 (endothelial cells). **C.** The percentage of infected cells that produced IFN differed significantly across primary human lung cell types. Each dot represents an experimental replicate. **D.** Representative smHCR images of influenza-infected human cell lines at 6 hpi. **E.** The percentage of infected cells that produced IFN+ differed significantly across cell types. Each dot represents an experimental replicate. Average n = 3969 (A549), 2640 (HDF), 1939 (HUVEC). **F.** The dependency of IFN-producing probability on the viral load for different primary human lung cell types at 8 hpi. Infected cells were sorted and binned based on viral load. r indicates Spearman’s rank correlation coefficient, and asterisks denote statistical significance. **B, C, E**: Error bars indicate mean +/- SEM; Asterisks denote statistical significance using one-way ANOVA test; n.s. denotes no significance.

### The endogenous RIG-I protein level controls the probability of pathway activation

Our single-cell analysis revealed remarkable heterogeneity in the probability of IFN production both within and across cell types, raising the question of what cellular factors drive this heterogeneity. By first examining the heterogeneity within a single cell type using influenza infection in A549 cells, we found that activation of the kinases immediate downstream of the viral sensing step, including IKK and TBK1, was limited to 3-5% of infected cells, and concurrently, only 1-4% of infected cells had nuclear translocation of the corresponding transcription factors, namely NF-κB and IRF3 (**Fig. 4A-C**). In contrast, >95% of cells with activated transcription factors produced IFN (**Fig. 4D, E**). These results suggest that the bottleneck of RIG-I pathway activation is the initial viral-sensing.

**Figure 4.**
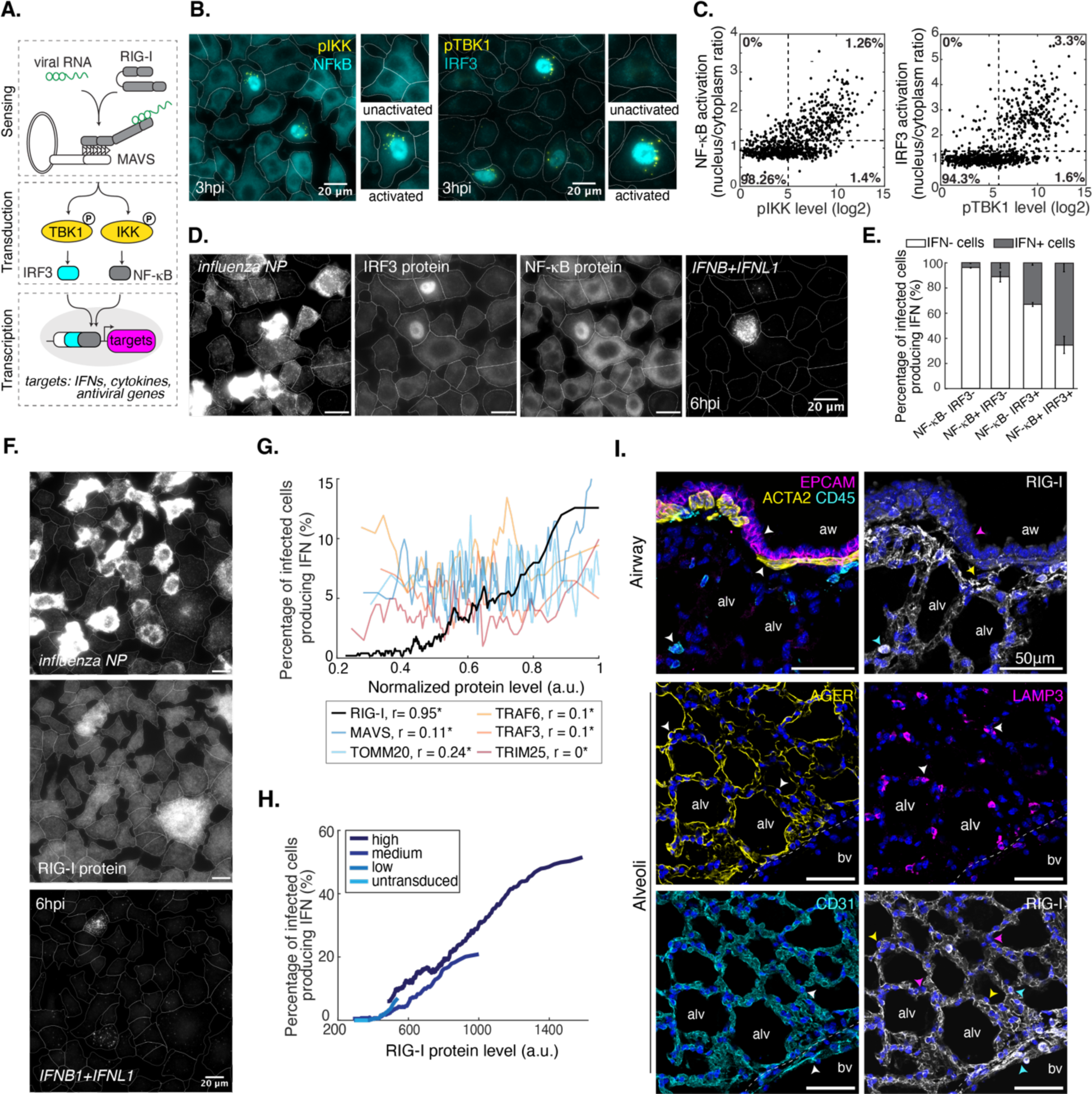
The endogenous RIG-I level controls the probability of pathway activation. **A.** Schematics of RIG-I pathway. Upon binding of viral RNA, activated RIG-I assembles the signalosome with the transmembrane adaptor mitochondrial antiviral signaling protein (MAVS), which phosphorylates and activates TANK-binding kinase 1 (TBK1) and IκB kinase (IKK). TBK1 and IKK then phosphorylate interferon-regulatory factor 3 (IRF3) and nuclear factor κB (NF-κB) transcription factors, leading to their nuclear translocation and transcription of *IFN*.^57–60^ ^B^. Representative immunofluorescence images of influenza-infected A549 cells co-stained for phosphorylated IKK (pIKK) and NF-κB (*left*) or phosphorylated TBK1 (pTBK1) and IRF3 (*right*) at 3 hpi. Insets: zoom-in on cells with inactivated and activated kinases. **C.** High efficiency of transcription factor activation in cells with activated kinases. n = 22,743 (NF-κB/pIKK), n = 14,138 (IRF3/pTBK1). **D.** Representative multiplexed smHCR (*NP* and *IFN)* and immunofluorescence (IRF3 and NF-κB) images of influenza-infected A549 cells at 6 hpi. **E.** High efficiency of *IFN* induction in cells with activated transcription factors. Error bars indicate mean +/- SEM, 3 replicates. **F.** Representative multiplexed smHCR (*NP* and *IFN)* and immunofluorescence (RIG-I) images of influenza-infected A549 cells at 6 hpi. **G.** The expression level of RIG-I, but not other pathway components, positively correlated with the probability of IFN production. For each protein, the expression level was normalized to its own maximum. Infected cells were sorted and binned based on the expression level of each protein to calculate the probability of IFN production. r indicates Spearman’s rank correlation coefficient, and asterisks denote statistical significance. **H.** Varying RIG-I protein level was sufficient for tuning the probability of IFN production. Lentiviral vector expressing RIG-I was titrated to achieve different levels of RIG-I reconstitution in *RIG-I-/-* A549 cells (see **Fig. S3F**). **I.** RIG-I protein level was high in immune and endothelial cells and low in epithelial cells in healthy adult mouse lungs (arrowheads). Tissues were labeled with antibodies against EPCAM (epithelia), ACTA2 (smooth muscle), CD45 (immune), CD31 (endothelia), LAMP3 (alveolar type 2), AGER (alveolar type 1), and mouse RIG-I. aw, airway; alv, alveolus; bv, blood vessel. See **Fig. S3G** for validation of mouse RIG-I antibody. All cells were treated with 10 μM Ruxolitnib in **B**-**H**.

Directly detecting activated RIG-I via imaging the assembly of RIG-I signalosome is challenging, due to the extreme rarity of such complexes ^30,31^. Protein concentrations are crucial parameters for complex assembly and previous studies suggested variability in the expression levels of RIG-I pathway components could contribute to the stochasticity of pathway activation ^17,19^. Therefore, we quantified the abundance of signalosome components and found that only the protein level of RIG-I highly correlated with the probability of cells producing IFN (**Fig. 4F, G, S3A**). Interestingly, RIG-I protein level was also more variable than other proteins measured, spanning ∼10 folds in uninfected A549 cells, which was not altered at 6 hpi (**Fig. S3B, C**).

To test the causal relationship between RIG-I level and the probability of IFN induction, we knocked out *RIG-I* in A549 cells and reconstituted RIG-I expression via lentiviral transduction. When infected with influenza, *RIG-I-/-* cells could be efficiently infected but did not produce any IFN, whereas cells with RIG-I reconstituted regained the capability of producing IFN (**Fig. S3D, E**). Titrating the doses of lentivirus resulted in different fractions of RIG-I+ cells and different levels per cell (**Fig. S3F**). Importantly, the probability of producing IFN still correlated well with the RIG-I protein level, confirming that RIG-I level is a critical tuning knob to regulate the probability of viral detection (**Fig. 4H**). We directly pinpointed the direct causal relationship between basal RIG-I level and the stochastic IFN production in lung epithelial cells.

Furthermore, we found highly variable RIG-I protein levels among different cell types in the mouse lung (**Fig. 4I, S3G**). The airway and alveolar epithelial cells expressed a low level of RIG-I, whereas a subset of the cells in the stromal compartment expressed a strikingly high level of RIG-I, contributing to the spatially graded probability of pathway activation. Interestingly, cultured human epithelial, fibroblast and endothelial cell lines that originated from different organs (e.g. lung, skin, and umbilical cord) retained the differences in RIG-I level that corresponded to their characteristic probability of pathway activation, suggesting the RIG-I level is a stable intrinsic feature of each cell family (**Fig. S3H, Fig. 3D, E**). Together, the basal level of RIG-I acts as a simple yet effective knob to control the probability of detecting infection, creating a stable spectrum of probability within a single cell type and across cell types.

### Tiered threat-sensing as a general principle for organizing barrier organ’s intrinsic immunity

Besides influenza viruses, the lung is exposed to a multitude of other pathogens and stimulants that are detected by other PRRs. We analyzed the expression levels of PRRs across cell types using scRNA-seq atlases of adult mouse and human lungs, as well as our own spatial transcriptomics data ^32–34^. The expression of many PRRs followed a similar trend as *RIG-I*, increasing from epithelial to endothelial cells (**Fig. 5A-C**). Interestingly, intracellular nucleic acid sensors (e.g. MDA5, TRL3, NOD1 and cGAS) exhibited a consistent trend, whereas PRRs on the plasma membrane (e.g. TLR2/4/5) had more variable patterns (**Fig. 5A-C, Fig. S4A, B**). Although the spatial expression patterns of membrane-bound PRRs were less clear, these PRRs have been shown to localize to the basolateral side of epithelial cells, making them less sensitive to pathogen-associated molecular patterns (PAMPs) unless the epithelial layer is breached ^35–39^. Therefore, lowering epithelial cells’ reactivity is a widely implemented design principle. Moreover, this differential expression pattern was unique to PRR expression and was not observed in other PRR pathway components, further confirming that the cell-to-cell variation originates at the receptor level (**Fig. S4C**).

**Figure 5.**
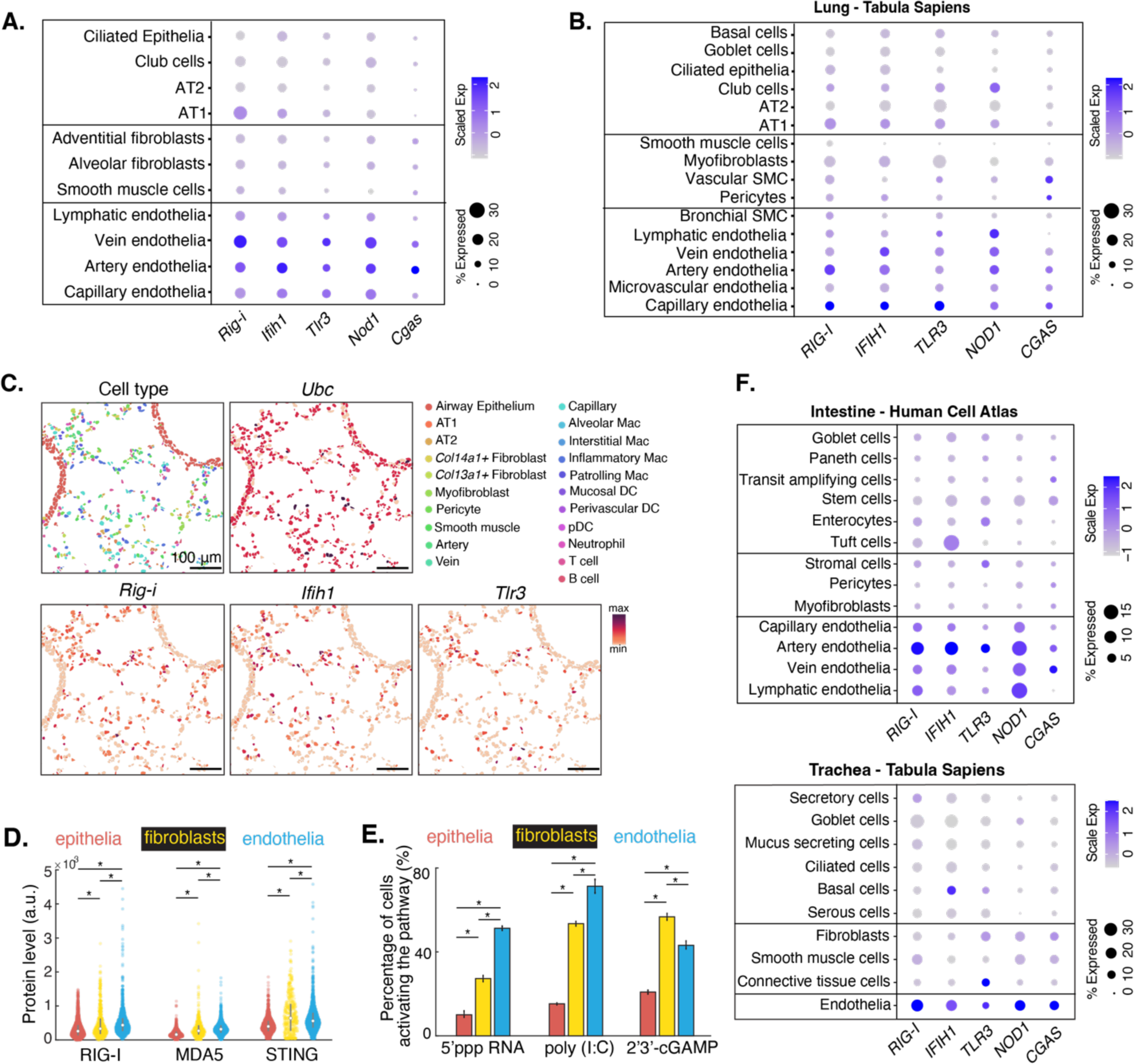
Tiered threat-sensing as a general principle for organizing barrier organs’ intrinsic immunity. **A.** Differential expression of nucleic acid-sensing PRRs among structural cell types in adult mouse lung using published scRNA-seq data ^34,61^. **B.** Same analysis as A using adult human lung data ^32^. SMC, smooth muscle cell. **C.** Expression of nucleic acid-sensing PRRs in adult mouse lung using STARmap PLUS data. *Ubc*, house-keeping gene control. **D.** Protein levels of nucleic acid-sensing PRRs were lowest in primary lung epithelial cells and highest in primary lung endothelial cells, quantified by immunofluorescence. Error bars, mean +/- SEM. **E.** The intrinsic probability of detecting synthetic PRR agonists among different primary lung cell types was generally correlated with the expression level of the corresponding receptor. Error bars indicate mean +/- SEM, 3 replicates. **F.** Expression levels of PRRs in adult human trachea and small intestine showed similar patterns as those in the lung ^32,62^. **D** & **E**: Asterisks denote statistical significance using one-way ANOVA tests.

Next, we asked whether the expression level of these PRRs correlated with the probability of pathway activation among different cell types. We first confirmed that the expression of PRRs in various human primary lung cell types showed a trend consistent with the transcriptomics data (**Fig. 5D**). We then challenged these cells by transfecting agonists specific to each PRR and found that epithelial cells activated PRR pathways at the lowest probability, in contrast to fibroblasts and endothelial cells, even though the transfection efficiency was comparable among different cell types (**Fig. 5E**, **Fig. S4D**) ^40–42^.

Furthermore, we observed a similar increase of PRR expression from the outermost to the innermost layers of the tissue in other barrier organs, e.g. the trachea and the intestine (**Fig. 5F, S4E**), which explains the previous observations on the low responsiveness to PRR ligands in intestinal epithelial cells ^43^. A similar down-regulation of PRRs in the outer barrier cells in the skin was observed, but the comparison was made among different layers of skin epithelial cells, whereas our observation of differential PRR expressions was among cell types from very different developmental origins ^44^. This raises an interesting question of how cells from unrelated lineages coordinate the expression levels of PRRs. In contrast, the tiered expression trend was not observed in non-barrier organs we analyzed, e.g. the ovary and breast (**Fig. S4F**). Although no functional tests were performed, these analyses suggest that tiered threat-sensing implemented by tissue structural cells is a general strategy of barrier organs for organizing their innate immunity.

### Increasing RIG-I level in lung epithelia exacerbates tissue damage under inflammatory challenge

Being the first cell type that encounters many insults, the low probability of activating PRRs in lung epithelial cells is surprising. We hypothesized that this low activation probability in epithelial cells might be a tradeoff between efficient self-defense and the need to avoid spurious immune activation to pathogen or pathogen-free stimuli that barrier cells frequently encounter. To test this hypothesis, we overexpressed RIG-I specifically in epithelial cells by delivering either a full-length mouse RIG-I or a truncated nonfunctional version (RIG-I-stop) directly into the trachea via lentiviruses (**Fig. 6A, S5A**). miR-142 binding sites in the 3’ UTR of the constructs represses transgene expression in hematopoietic cells and reduces transgene toxicity ^45,46^. Successful transduction and integration of the constructs was monitored by the co-expression of mScarlet, which showed a consistent transduction efficiency of ∼2% among airway epithelial and alveolar cell types, with minimal expression in other cell types (**Fig. 6B, S5B**). mScarlet+ cells were predominantly epithelial cells (∼85%), distributed equally between airway and alveolar cell types, even up to 8 weeks post transduction (**Fig. 6C, S5C**). We further confirmed that the inflammatory response caused by intratracheal instillation of viral vectors resolved after two weeks, as measured by the abundance of immune cells within the tissue and in bronchoalveolar lavage (BAL) (**Fig. 6D, S5D**).

**Figure 6.**
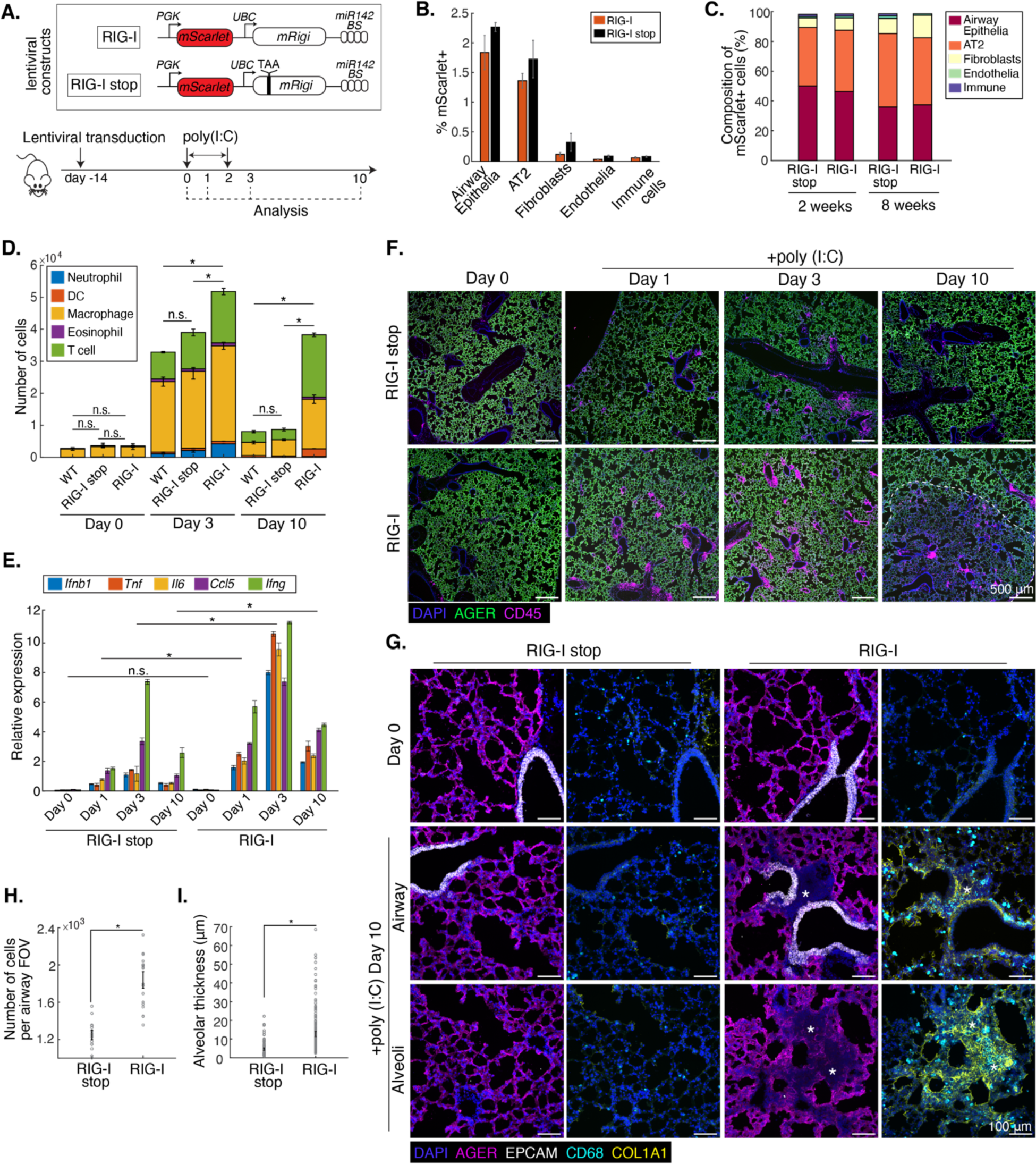
Increasing RIG-I level in lung epithelia exacerbates tissue damage under inflammatory challenge. **A.** Schematics of experimental design. Lentivirus expressing RIG-I or RIG-I stop (an early stop codon at amino acid 49) was transduced into the lung of wild-type mice. Two weeks later, mice were challenged daily with 25 μg low molecular weight poly(I:C) intra-nasally for three consecutive days (day 0-2). The lung was analyzed on different days; analysis on day 0 was done prior to poly(I:C) dosing. BS, binding sites. **B.** Quantification of transgene-expression efficiency among different cell types in the lung at 2 weeks post lentiviral transduction as indicated by mScarlet+ on flow cytometry. n = 5 mice per group. **C.** The majority of mScarlet+ cells were airway and alveolar epithelial cells at 2 weeks and 8 weeks post lentiviral transduction. **D.** Immune cell infiltration in bronchoalveolar lavage (BAL) upon poly(I:C) injection. n = 2-3 mice per group per time point. **E.** Levels of cytokines and chemokines upon poly(I:C) injection, quantified by RT-qPCR. Each cytokine/chemokine was compared separately, matching time points for two groups. n = 2-3 mice per group per timepoint. **F.** Immunofluorescence staining of lung sections before and after poly(I:C) injection. Tissue damage and immune-cell infiltration were analyzed with antibodies against AGER (alveolar type 1) and CD45 (immune cells). Dotted line marks the damaged region. **G.** Immunofluorescence staining of lung sections at day 10 after poly(I:C) injection. Damage and repair were analyzed with antibodies against EPCAM (pan-epithelia), AGER (alveolar type 1), CD68 (macrophage), and COL1A1 (collagen). Asterisks mark damaged regions. **H.** Increase in airway cellularity in mice overexpressing RIG-I. Cellularity was measured by the number of nuclei per airway field of view (FOV). n = 8 sections across 2-3 mice per group. **I.** Increase in alveolar thickness in mice overexpressing RIG-I. Alveolar thickness was measured using length of lines connecting two adjacent alveoli. n = 8 sections across 2-3 mice per group. **B**, **D**, **E**, **H**, **I**: Error bars indicate mean +/- SEM. Asterisks denote statistical significance and n.s. denotes no significance using one-way ANOVA tests.

To induce inflammation in the lung, we administered low molecular weight poly(I:C) intra-nasally, which transfected 30% of epithelial cells, accounting for 60% of the overall uptake (**Fig. S6A, B**). The dose caused mild immune infiltration in the wild-type lung (**Fig. S6C**). Under the same poly(I:C) challenge, RIG-I mice had much stronger RIG-I pathway activation and IFN signaling, compared to RIG-I-stop mice (**Fig. S6D**). Animals in both cohorts engaged in a global inflammatory response; however, RIG-I mice expressed cytokines and chemokines at a higher amplitude compared to RIG-I-stop mice (**Fig. 6E**). Consistently, we also observed a significantly higher number of immune cells in BAL in RIG-I mice at day 3 (**Fig. 6D**). Overall, having a higher level of RIG-I in epithelial cells leads to more potent immune responses at the onset of inflammatory challenge.

Although strong and early immune responses can clear pathogens quickly, high magnitude of immune activation could impact resolution of inflammation and subsequent tissue repair. We found that RIG-I mice sustained a high level of cytokines and chemokines in the tissue and infiltration of immune cells in BAL at 10 days post-poly(I:C) treatment. In contrast, inflammation in RIG-I-stop mice significantly resolved at day 10 (**Fig. 6D, E**). Consequently, RIG-I mice suffered from a significant damage to the alveolar structure and a prolonged infiltration of immune cells in the tissue (**Fig. 6F, S6E, F)**. Furthermore, in these inflamed regions, we observed an enriched deposition of Collagen 1, a significant increase in airway cellularity, and a dramatic thickening of the alveolar wall, suggesting impairment in respiratory function and a potential onset of fibrosis in RIG-I mice post pathogen-free inflammation (**Fig. 6G-I**) ^47^. Together, these results provide strong evidence that keeping the sensitivity low in epithelial cells can be beneficial for the host, by preventing spurious activation of the immune responses when facing mild, non-pathogenic insults.

## Discussion

By asking how organs evaluate the level of threats and adjust the appropriate immune responses, we discovered an unexpected tissue-scale strategy to balance host protection with collateral tissue damage at the onset of threats. This strategy emerges from the differential intrinsic capability of sensing threats among different cell types that are spatially organized into distinct layers. Specifically, the outermost epithelial cells have the lowest probability of detecting threats, and cells in the stromal compartment, especially endothelial cells, have the highest probability of detecting threats. This spatially tiered threat-sensing design allows the tissue to evaluate the degree of infection and adjust the fraction of cells producing cytokines accordingly. Cytokines recruit immune cells to infected tissues, but massive infiltration of immune cells can cause tissue damage ^48,49^. By adapting the fraction of responsive cells to match infection severity, the total amount of cytokines can be quantitatively regulated to balance the benefits and the costs of immune activation. Importantly, the high sensitivity of cells in the stroma acts as a “fail-safe” mechanism that rapidly ramps up the immune response when the epithelial barrier is breached and the threat penetrates deeper into the tissue, to prevent infection from spreading further into the circulation and to other vital organs. Together, the collaboration among different cell types represents a novel way to collectively tune the magnitude of initial immune response during infection.

The low pathogen-sensing capability in epithelial cells appears sub-optimal for protecting hosts from infection, but we demonstrated that it can benefit the host when facing non-infectious challenges. It could contribute to the tolerance of commensal or non-pathogenic microbes colonizing barrier organs. Barrier sites are also subjected to frequent low-grade mechanical perturbations that trigger the release of danger-associated molecular patterns (DAMPs) ^50,51^. Because PRRs lack specificity in distinguishing between DAMPs and PAMPs, the low sensitivity among epithelial cells might also help minimize spurious immune activation to DAMPs ^52^.

Although the influenza strain used in this study has wide tropism in mice ^29,53^, whereas most pathogens have more limited tropism ^54,55^, this spatially tiered threat-sensing design can be useful for tuning the initial immune responses regardless of the pathogen’s tropism in the lung. Pathogens that only infect epithelial cells pose less threat to the host, whereas pathogens that quickly infect endothelial cells are more likely to cause damage to the host. Therefore, the different probabilities of activating the PRR pathways among different types of cells still allow the host to regulate the magnitude of immune responses based on the severity of the initial infection.

In conclusion, we revealed how a tiered threat-sensing mechanism emerges in the lung and its potential functions, and provided evidence that this tiered design of innate immunity might be universal across barrier organs. The differential expression of PRRs among epithelial cells, fibroblasts, and endothelial cells is observed in *in vivo* human and mouse tissues, in primary cells from the lung, and in immortalized human cells from various organs. Despite the differences in their origin and culture conditions *in vitro*, the characteristic PRR expression levels and detection sensitivity are well preserved for each cell class. This suggests a robust intrinsic regulation of PRRs associated with the identity of the cell class. When and how these different cell classes gain their signatures of PRR levels across multiple PRRs, and how these universal functional states tie with cell identity remain to be revealed.

### Limitations

This study used naive pathogen-free mice, which is potentially the reason why a low proportion of tissue-resident immune cells was detected. It would be interesting to measure the contribution of tissue-resident immune cells to the initial detection of influenza virus in mice with prior pathogen exposure. We used mRNA levels as a proxy for viral load and IFN production, because the actual RNA species detected by RIG-I still remains elusive. Although defective viral RNA has recently been implicated as the agonist for RIG-I, it would be impossible to probe for with our current methods at the single-cell level ^56^. A measurement of IFN protein level would be more accurate than mRNA; however, given the fast secretion of cytokines upon synthesis, it would be difficult to quantify the intracellular protein level *in vivo* with single-cell resolution. Lastly, when controlling for the transfection efficiency in different primary human lung cells and *in vivo*, a fluorescent non-immunogenic small RNA was cotransfected with the agonists of interest and used as a proxy for the level of agonists per cell, due to the lack of access to agonists directly conjugated to a fluorophore. Even with such reagents, it is still difficult to quantify the amount of agonists that successfully escape the endosome, bind PRRs, and trigger the responses.

## Resource availability

### Lead contact

Requests for further information and reagents should be directed to the lead contact, Pulin Li, at.

### Materials availability

Plasmids and cell lines generated in this study will be available upon request.

### Data and code availability

The STARmap PLUS sequencing data and codes for STARmap analyses are available at https://github.com/Pulin-Li-Lab/Tiered-immunity-2025.

## Acknowledgements

We would like to thank Xu Zhou, Sebastian Lourido, Harikesh S. Wong, and Jordi Garcia-Ojalvo for valuable advice and support, Jianzhu Chen for providing critical reagents, Mark Greenwood, Daniel Y. Zhu for advice on the manuscript, in addition to members of the Li Lab for discussion and feedback through all stages of the project. We would also like to thank the flow cytometry core facilities (Whitehead Institute and Koch Institute), the W.M.Keck Microscopy Facility (Whitehead Institute), and the Hope Babette Tang Histology Facility (Koch Institute) for providing access to their instruments. This work was supported by National Institute of Health grants DP2HD108777 (P.L.) and 1DP2GM146245 (X.W.), Searle Scholars Program (X.W.), Thomas D. and Virginia W. Cabot Professorship (X.W.), Edward Scolnick Professorship (X.W.), Merkin Institute Fellowship (X.W.), National Cancer Institute grant R35-CA274464 (T.J.), David H. Koch Graduate Fellowship (S.L.S), MIT School of Science Fellowship in Cancer (S.L.S) and MathWorks Predoctoral Fellowship (D.H.N.). T.J. is a Daniel K. Ludwig Scholar.

## Author contributions

Conceptualization: D.H.N., P.L.; Investigation: D.H.N., J.T., S.L.S., C.K.W.; Resources: X.W., T.J.; Funding acquisition: X.W., T.J., P.L.; Writing: D.H.N., P.L.; Supervision: P.L.

## Supplemental information

Materials and Methods

Supplementary Figures 1–6

Supplementary Tables 1-2

## Materials and Methods

### Mice

All experimental procedures involving the use of mice were performed in compliance with the *Guide for the Care and Use of Laboratory Animals*. All experimental procedures were approved by the Massachusetts Institute of Technology (MIT) Committee on Animal Care (Protocol #2403-000-640). Wild-type, 6-12 week-old female mice in BALB/c background (for influenza infection and *in situ* sequencing) and C57BL/6J background (for lentiviral transduction and inflammatory challenge) were purchased from Jackson Laboratories (Bar Harbor, Maine) and were between 8 and 14 weeks of age at the time of influenza infection or lentiviral infection. All mice were housed as groups of up to five animals per cage when possible and supplied with extra enrichment if singly housed. All mice were fed with regular rodent’s chow and sterilized water.

### Cell lines and culture

Human lung epithelial carcinoma line (A549) (ATCC CCL-185), Madin Darby canine kidney cell line MDCK (gifted by Dr. Jianzhu Chen, Koch Institute/MIT), and primary human dermal fibroblast cells (gifted by Dr. Sebastian Lourido, Whitehead Institute/MIT) were maintained in DMEM supplemented with 10% fetal bovine serum (Takara Bio, 631368), 1% Pen/Strep/L-Glutamine. Primary human umbilical vein endothelial cell line (HUVEC) was purchased from the Preclinical Modeling core facility at the Swanson Biotechnology Center, Koch Institute, MIT and maintained in VascuLife® VEGF Endothelial Medium Complete Kit (LL-0003). Primary human bronchial epithelial cells (H-6033B), lung fibroblasts (H-6013), and pulmonary vein cells (H-6060) were purchased from Cell Biologics, plated on 0.1% gelatin-coated plates and maintained in the corresponding complete media per instruction of the supplier.

### Virus

For *in vivo* influenza infection in mice, influenza A virus A/WSN/33 (H1N1) expressing NanoLuc appended to the C terminus of PA (PASTN)^63^ was obtained through BEI Resources, NIAID, NIH (NR-49383). For influenza infection in cultured cells, influenza A virus A/WSN/33 (H1N1) was gifted by Dr. Jianzhu Chen (Koch Institute/MIT).

### Chemical agonists and antagonists

Pan-IFN response was blocked by the addition of 10 μM Ruxolitnib (Sellekchem, S1378) into culture medium during viral infection. To activate PRR pathways in cultured primary cells, the following agonists were used: 250 ng/mL 5’ppp-hairpin RNA (Invivogen, tlrl-hprna), 100 ng/mL high molecular weight poly(I:C) (Invivogen, tlrl-pic), and 50 μg/mL of 2’3’-cGAMP (Invivogen, tlrl-nacga23-02). 5’ppp-hairpin RNA and poly(I:C) were transfected using Lipofectamine RNAiMAX (Invitrogen, 13778150)

### *In vivo* influenza infection

Mice were anesthetized with 3-5% isoflurane. 40 μL of influenza virus (a total of 10^4^ PFUs) was delivered drop-wise intranasally per mouse using P20 pipette. Mice were weighed daily, assessed for morbidity according to guidelines set by the MIT Division of Comparative Medicine, and were humanely sacrificed prior to losing >20% of the initial body weight.

### Influenza A virus production and titer

Influenza A virus was propagated by infecting MDCK cells at an MOI of 0.1 in MEM media supplemented with 0.15% sodium bicarbonate, 0.01M HEPES, 0.2% bovine serum albumin (BSA), and 1% Pen/Strep/Glutamine. Cells were incubated for 36 hr prior to harvesting virus supernatant. The supernatant was centrifuged at 500xg for 5 min at 4℃ to pellet cellular material. Clarified supernatant was aliquoted into 500 μL - 1mL each, stored at −80℃, and titered using plaque assay.

Plaque assay was performed by infecting MDCK with 10-fold serial dilution of the viral stock. After 1 hr incubation, virus was removed; 1 X virus media in 0.6% low-melting agarose was added and let solidified at room temperature for 10 min before incubating at 37℃ for 48 hr. To determine the number of plaque-forming units (PFU) in the viral stock, immunofluorescence staining of NP protein on infected MDCK was performed, and the number of foci was counted manually using EVOS M7000.

### Mouse lung tissue collection for smHCR, STARmap PLUS, and immunofluorescence staining

Mice were euthanized with 5% CO_2_. After euthanasia, mice were perfused by intracardiac injection of 1-20 mL PBS into the right ventricle until the lung appeared opaque. After perfusion, the lung was inflated with 1.5 mL of 50% OCT: 50% PBS through intratracheal instillation. Following inflation, the trachea was quickly clamped with a hemostat. Lung was removed en-bloc, embedded in 100% OCT, flash frozen using liquid nitrogen, and stored at −80℃.

For tissue sectioning, mouse lungs were transferred to cryostat (Leica CM3050 S) and cut into 20 μm thick slices from proximal to distal at −20℃. The slices were attached to each well of glass-bottom 24-well plates pretreated by methacryloxypropyltrimethoxysilane (bind-silane) and poly-D-lysine (for STARmap PLUS) and poly-D-lysine only (for others). The lung slices were fixed with 4% PFA in 1 X PBS buffer at room temperature for 15 min, then permeabilized with cold methanol and placed at −20℃ for 30 min before hybridization/immunostaining.

### STARmap PLUS

Due to the spatial localization of infection foci at 36 hpi and the sparsity of infection foci within the lung tissue, we reasoned that, given an averaged focus size of 50-100μm, adjacent sections should show similar infection patterns. Therefore, we sectioned the infected lung tissues into two 24-well plates where the two adjacent sections were placed into separate plates. To determine which sections have infection foci, we performed immunofluorescence staining of influenza NP on all sections from one plate, examined infection patterns under Leica Stellaris 8, and proceeded with the STARmap PLUS protocol on the corresponding positive sections of the second plate.

STARmap PLUS protocol was performed as previously reported^63^. Briefly, after lung tissue sections were fixed and permeabilized, the intracellular mRNAs were targeted by a pair of SNAIL (specific amplification of nucleic acids via intramolecular ligation) probes (IDT). The probes were then enzymatically ligated and amplified to generate amine-modified cDNA amplicons *in situ*. Subsequently, the same tissue sections were labeled with rabbit anti-influenza NP polyclonal primary antibody (Invitrogen, PA5-32242) at 1:500 dilution. Next, tissue sections with amine-modified cDNA amplicons, proteins and primary antibodies were modified by methacrylic acid N-hydroxysuccinimide ester (MA-NHS) and copolymerized with acrylamide and bis-acrylamide to generate a hydrogel-tissue hybrid that fixes the locations of biomolecules for *in situ* mapping. Lastly, fluorescent protein staining via goat anti-rabbit secondary antibody (Invitrogen, A-21244) was performed to visualize infection foci.

Fields-of-view for sequencing were chosen based on influenza NP staining signal to include both uninfected and infected regions from various anatomical locations. The fluorescent signal of the secondary antibody was quenched via a brief proteinase digestion and photobleaching prior to *in situ* sequencing. Images were acquired using Leica Stellaris 8 confocal microscopy (Leica LAS-X microscope imaging software) with a 405 nm diode, a white light laser, and a 40x oil immersion objective (NA 1.3). The laser lines we used for SEDAL sequencing are 488 nm, 546 nm, 594 nm and 647 nm. We imaged with a voxel size of 142 nm (*x* axis) × 142 nm (*y* axis) × 350 nm (*z* axis) and 45 z slices (15 µm in total). Five cycles of imaging were performed to detect 237 genes and DAPI (4’,6-diamidino-2-phenylindole) signal was acquired in the first cycle.

### Single-molecule HCR RNA-FISH (smHCR)

*In situ* HCR v3.0 was performed according to the manufacturer’s protocol (Molecular Instruments)^64^. Probes were designed in-house, and amplifiers with buffers were supplied by Molecular Instruments.

#### For fresh-frozen tissue sections

Following fixation and methanol permeabilization, lung slices were removed from −20℃ to room temperature for 5 min. The samples were washed once with quenching buffer (0.1M glycine in 1X PBS with 0.1% Tween-20, 0.1U/μL SUPERase-In RNase inhibitor) and once with 1X PBS with 0.1% Tween-20, 0.1U/μL SUPERase-In RNase inhibitor (PBSTR). After washing, the samples were incubated with 250 μL hybridization buffer with 1 μM per probe in a 37℃ humified oven with parafilm wrapping for 16-24 hr. Probes were removed with a formamide-containing wash buffer. After washing, the slices were treated with 3.75 pmol of the amplifiers in 125 μL of amplification buffer at room temperature for at most 90 min. The amplifiers consist of a pair of hairpins conjugated to the following fluorophores: Alexa 488, 546, 594, and 647. Excess hairpins were then washed with 5X SSCT (sodium chloride sodium citrate with 0.1% Tween-20). Nuclei were counterstained with DAPI.

#### For cell culture

Samples were washed twice with 1X DPBS, fixed with 4% PFA for 10 min at room temperature, washed twice with DPBS to remove PFA, and permeabilized with 100% methanol for 10 min at −20℃. Samples were washed twice with 2X SSC and incubated with 250 μL hybridization buffer with 1 μM per probe in a 37℃ humified oven with parafilm wrapping for 16-24 hr. Probes were removed with a formamide-containing wash buffer. After washing, the slices were treated with 3.75 pmol of the amplifiers in 125 μL of amplification buffer at room temperature for 45 min. The amplifiers consist of a pair of hairpins conjugated to the following fluorophores, Alexa 488, 546, 594, and 647. Excess hairpins were then washed with 5X SSCT (sodium chloride sodium citrate with 0.1% Tween-20). Nuclei were counterstained with DAPI.

Images were collected under Zeiss 710 LSM laser scanning confocal with 40x oil objective (NA 1.3) and Nikon Ti2 with a 60x oil objective (NA 1.4) and a Zyla CMOS detector. Samples were imaged on a z step of 15 z-planes spanning 15 μm for lung tissue section and 7.5 μm for cell culture.

### Immunofluorescence staining

For both tissue sections and cell culture, following fixation and methanol permeabilization, samples were incubated in a blocking buffer containing 1X PBS with 0.1% Tween-20 (PBST) and 10% normal goat serum or 1% BSA for 30 min at room temperature. After the blocking buffer is removed, samples were incubated with primary antibodies for 2 hr at room temperature or overnight at 4℃. Samples were then washed three times with PBST. Secondary antibodies were treated for 1 hr at room temperature. Samples were washed three times in 1X PBST at room temperature and counterstained with DAPI.

### Multiplex single-molecule HCR RNA-FISH and immunofluorescence staining

Following fixation and methanol permeabilization, lung slices were removed from −20℃ to room temperature for 5 min. The samples were washed once with a quenching buffer and once with PBSTR. After washing, samples were incubated in a blocking buffer containing 0.1% Ultrapure BSA (Invitrogen) in PBSTR for 30 min at room temperature. Next, primary antibodies diluted in blocking buffer were added to the samples and let incubate for 90 min at room temperature. Samples were washed three times with PBSTR. Secondary antibodies were treated for 30-45 min at room temperature. Samples were washed three times with PBSTR. Samples were post-fixed with 4% PFA for 10 min, washed twice with 2x SSC and continued with smHCR protocol.

### Western blotting

Western blotting was performed using standard procedures. Briefly, ∼10^6^ cells were trypsinized in 0.25% trypsin, then trypsin was neutralized with tissue culture medium, and cells were centrifuged at 500xg for 5 min. Cells were washed with DPBS, then lysed on ice for 10 min in 100 µL Tris-Triton lysis buffer (10 mM Tris pH 7.4; 100 mM NaCl; 1 mM EDTA; 1 mM EGTA; 1% Triton X-100; 10% glycerol; 0.1% SDS; 0.5% deoxycholate) with protease inhibitors (cOmplete Mini, Roche). Extracts were mixed with reducing agent (Life Technologies, NP0004) and LDS sample buffer (Life Technologies, NP0007), then heated at 95°C for 10 min. Extracts were chilled on ice, then centrifuged for 2 min at 15,000×g, and 10 µL soluble protein was loaded into a precast NuPAGE 3 to 8% Tris-Acetate gel (Invitrogen). Proteins were transferred using an iBlot2 (Invitrogen) at 25 V for 10 min. The membrane was blocked with Licor TBS blocking buffer for 30 min at room temperature, then detected overnight at 4°C with mouse anti-RIG-I monoclonal primary antibody (clone Alme-1, AdipoGen Life Sciences) at a 1:1000 dilution. As a loading control, the membrane was labeled with mouse anti-alpha-tubulin primary antibody (Cell Signaling #3873) at a 1:1000 dilution. LiCor goat anti-mouse antibodies were used to detect primary antibodies, and membranes were imaged using a Licor Odyssey Clx fluorescence scanner.

### Generation of *RIG-I-/-* A549 cell line

Wild-type A549 cells were transiently transfected with a Cas9-T2A-mCherry plasmid, encoding an sgRNA targeting the sequence GAAAAACAACAAGGGCCCAA. Transfected cells were sorted by FACS at 24 hr after transfection for mCherry(+) cells. Sorted cells were allowed to recover for 3-5 days and sorted again to generate single-cell clones of *RIG-I^−/−^* cells. Clones were screened by PCR & Sanger sequencing, and knockouts were validated by immunofluorescence staining and western blotting.

### Lentiviral constructs

RIG-I was reconstituted in RIG-I-/-A549 cells using plasmid pTRIP-SFFV-mtagBFP-2A RIGI WT (gifted by Agnel Sfeir, Addgene plasmid # 167289). RIG-I overexpression in mouse lung was accomplished by lentiviral constructs (LV-PGK-mScarlet-UBC-mRigi-mir142BS for full-length RIG-I protein, LV-PGK-mScarlet-UBC-mRigi_stop-mir142BS for truncated RIG-I protein). Constructs were generated using Gibson assembly. Lentiviral backbone carrying mScarlet and mir142BS was gifted by Dr. Tyler Jacks (Koch Institute, MIT). UBC promoter and mouse RIG-I fragments were amplified from the following plasmids: yWL229 MCP-GFP-eRF3 (gifted by Robert Singer, Addgene plasmid # 197053) and pLenti6.2_3xFLAG_V5_ mmDdx58_V5 (gifted by Rizwan Haq, Addgene plasmid # 160108). Lentiviral construct carrying Rigi_stop was generated by cutting the lentiviral construct carrying full-length RIG-I with restriction enzymes XhoI and NheI to introduce early stop codon at amino acid 49.

### Lentiviral production for *in vivo* instillation

Lentivirus was produced by transfection of LentiX-293T viral packaging cells in 15-cm plates with lentiviral constructs (10μg), VSV-G (2.5μg), and psPAX2 (7.5μg) viral packaging plasmids, and polyethylenimine (PEI, 60 μL). Lentiviral supernatant was harvested at 48 hr and 72 hr post-transfection, filtered through a 0.45 μm PVDF filter, and concentrated by ultracentrifugation at 25,000 RPM for 2 hr at 4°C. Viral titers were determined by measuring mScarlet expression in GreenGo 3TZ cells.

### Lentiviral transduction in mouse lung

Mice were anesthetized with 3-5% isoflurane. The airways were conditioned with 25 μL of 0.1% lysophosphatidylcholine (LPC, Sigma-Aldrich, L4129) One hour later, 50 μL of 6-8×10^5^ transduction units of lentivirus containing mScarlet-mRigi and mScarlet-mRigi-stop was delivered via intra-tracheal injection. Immediately prior to injection, all viral supernatants were mixed with LentiBOOST (SIRION Biotech) to improve transduction efficiency.

### *In vivo* injection of poly(I:C)

25 μg of low molecular weight poly(I:C) (Invivogen, tlrl-picw) was injected intranasally. Briefly, 25 μg poly(I:C) and 3 μL jetPEI (Polyplus Transfection) were diluted and mixed with 5% glucose solution to a total of 40 μL injection solution per mouse. After 15 min of incubation at room temperature, the complex was injected drop-wise intranasally using P20 pipette. Mice were anesthetized with 3-5% isoflurane for the injection. To quantify the efficiency of injected RNA getting into cells, 25 μg fluorescently labeled tracrRNA-647 complexed in jetPEI was injected intra-nasally into wild-type mice. Lung digest was collected at 16 hr post transfection and analyzed by flow cytometry.

### Mouse lung tissue dissociation for flow cytometry

After euthanasia, mouse lung tissue was collected in 5 mL HBSS supplemented with 5U/mL dispase II (STEMCELL technology, 07913), 125 U/mL collagenase IV (Worthington Biochemical, CLS-4), and 40 U/mL DNaseI (Sigma-Aldrich, 10104159001). The tissue was manually dissociated by mincing with scissors and pipetting. Dissociated tissues were incubated at 37°C for 30-45 min on a shaker table. Cell suspensions were centrifuged at 1700 RPM for 3 min and red blood cells were lysed using ACK lysis buffer (Gibco, A1049201). Single-cell suspension was filtered with 70 μm cell strainer and stained with conjugated primary antibodies (EPCAM-BV711, CD31-FITC, CD45-APC, MHCII-PacBlue, CD11c-PE-Cy7). Samples were analyzed on a BD Biosciences Aria 2.

### RNA extraction and quantification

Tissue samples were briefly dissociated, and RNA was extracted using RNA Miniprep kit (Zymo Research). cDNA was synthesized using MMLV reverse transcriptase (Biorad) and oligo(dT) primers. RT-qPCR reactions were performed on the Applied Biosystems QuantStudio3 Real-Time PCR System (ThermoFisher) using PowerUp SYBR Green Master Mix (ThermoFisher). Fold induction in RT-qPCR analysis was calculated as (coef^-ddCt), where dCt values were normalized to wild-type controls and coef was determined using standard curves. ΔCt was calculated by subtracting *Actb* Ct values from Ct values of genes of interest.

### BAL collection and analysis

BAL was collected by cannulation of the trachea, followed by instillation and aspiration of 1mL PBS. BAL cells and fluid were separated by centrifugation at 500xg for 5 min. BAL cells were stained with conjugated primary antibodies (MHCII-PacBlue, SIGLECF-Alexa488, LY6G-Alexa594, CD11b-APC, CD11c-PE-Cy7 CD3-BV711) and analyzed on a BD Biosciences Aria 2.

### Quantification and statistical analysis

All of the image processing steps and quantification were implemented using MATLAB R2021b, open-source packages in Python 3.10, and ImageJ 2.14.

### smHCR and IF staining image analysis

#### Single-cell segmentation

##### Cell culture

Overlay of nuclei by DAPI staining and cytoplasm by protein staining or *ACTB* RNA was generated by max-z projection. Cell bodies were identified by applying a pretrained 2D machine learning ‘cyto2’ model from Cellpose ^65^.

##### Lung tissue section

Nuclei were automatically identified by applying a pretrained 2D machine learning ‘nuclei’ model from Cellpose ^65^.

Segmentation overlay: The output file from Cellpose, ‘_cp.outlines.txt’, was loaded into ImageJ using ‘imagej_roi_converter.py’. The segmentation masks appeared as ROIs in ROI manager. *smHCR quantification*

To identify and count the spots corresponding to individual mRNA molecules, we developed an in-house MATLAB program where a Laplacian of Gaussian filter (sigma = 0.2) was applied to the input images to enhance the spots. To identify spots, the program applies an intensity threshold using Otsu thresholding and an area threshold.

#### Protein abundance quantification

Pixel intensity was background subtracted prior to analyses. Cell segmentation masks were loaded into MATLAB. Single-cell quantification of protein abundance was obtained using function regionprops with “MeanIntensity”.

#### Statistical analysis

*In situ* smHCR and IF images were analyzed using in-house MATLAB scripts for quantification and analysis. Mean, standard error of the mean, one-way ANOVA, Fisher’s exact test, Spearman correlation, and Pearson correlation were calculated using MATLAB. n.s. = no significance, * denotes significance (p < 0.05)

### STARmap PLUS image processing and analysis

#### Image processing

STARmap PLUS data analysis was performed similarly as described previously ^28,66,67^. Briefly, image deconvolution was performed with Huygens Essential version 21.04 (Scientific Volume Imaging, The Netherlands, http://svi.nl), using the CMLE algorithm, with SNR:10 and 10 iterations. Image registration, spot calling, and barcode filtering were performed as previously described. Cell segmentation was done using the sum z-projection of DAPI signal with watershed and dilation. NP staining was segmented and overlaid with the cell segmentation mask to identify NP-positive cells.

#### Cell type classification

Count matrices of regions of interest were loaded into Seurat v4 and concatenated. Counts were then divided by total counts and multiplied by a factor of 10000, followed by log transformation, all implemented in Seurat’s default setting. A set of 78 marker genes that are differentially expressed across cell types was curated to identify major cell types in adult mouse lung ^7,9^. Highly variable genes of the gene expression matrix were set to the chosen 78 marker genes. A PCA was applied to reduce the dimensionality of the cellular expression matrix. The top 25 PCs were used to compute a kNN graph of the observations. The Louvain algorithm with a resolution of 0.5 was applied to detect cell clusters. Clusters were annotated based on their top representative markers (a subset of markers shown in **Fig. S2B**).

#### Neighborhood analysis

A ‘window’ was captured consisting of the 14 nearest uninfected neighboring cells and a center as measured by Euclidean distance between X/Y coordinates. Cells with more than one count of influenza *NP* transcript are considered infected, otherwise uninfected. For uninfected neighborhoods, both the center cell and neighboring cells are uninfected. For infected neighborhoods, the center cell is infected and neighboring cells can be either infected or uninfected. For foci, only infected cells were considered. Infected cells which were close in distance were likely to belong to the same infection focus. Therefore, we applied density-based clustering (DBSCAN, implemented in scikit-learn, python 3.10) on infected cells’ X/Y coordinates to identify foci within each section.

#### Statistical analysis

To test the significance of cell type enrichment in different types of neighborhoods, Fisher exact test was used for each cell type and Benjamini-Hochberg correction was performed for multiple testing. n.s. = no significance, * = significance (p < 0.05). To test whether there is a significant difference in the probability of IFN induction within and among cell type families, Kruskal-Wallis test was used. For within cell type family tests, the probability of IFN induction of each cell type is compared to other cell types. For across cell type family tests, the probability of IFN production of cell types within the same cell family was pooled together, and Kruskal-Wallis test was performed on main cell families: epithelia, fibroblasts, endothelia, and immune. n.s. = no significance, * = significance (p < 0.05)

### Single-cell RNA-seq analysis

Datasets were loaded into Seurat v4, filtered, and normalized using standard Seurat’s pipeline. Differentially expressed PRR genes were identified by performing Wilcoxon rank-sum test using the ‘FindAllMarkers’ function in Seurat and setting features to a set of PRR genes. Significant genes have log fold-change > 0.1 with adjusted p-value < 0.05 and are expressed in at least 1% of the cell population.

---

**Figure S1.**
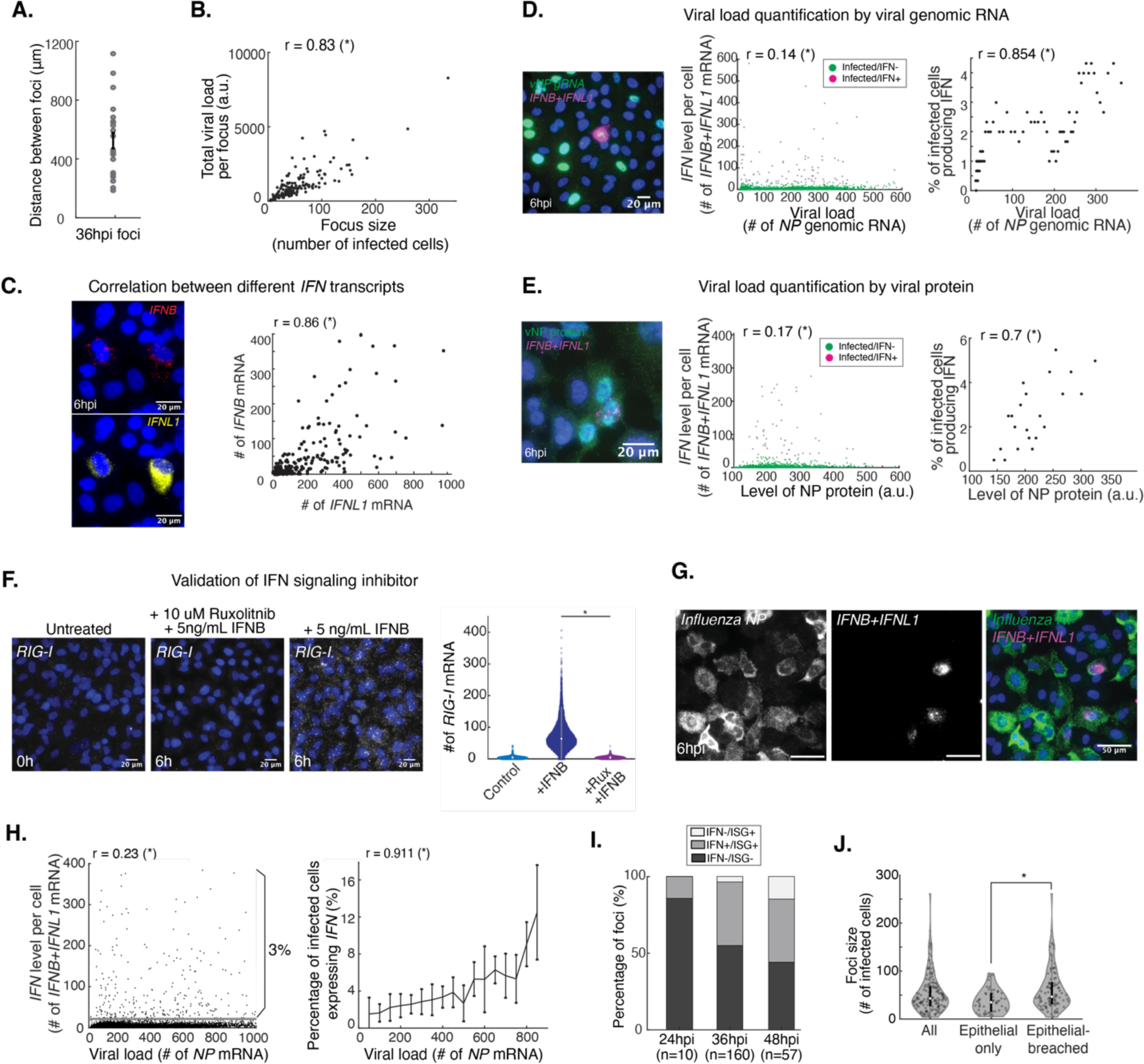
Probabilistic control of IFN production is an intrinsic property of infected cells (related to Figure 1). **A.** Quantification of 2-D distance between foci in a single 20 μm lung section. **B.** Focus size in the lung correlated with the total viral load. n = 160 plaques. **C.** Correlation between human *IFNB1* and human *IFNL1* mRNA at the single-cell level in human lung A549 cells, measured by smHCR. **D.** The percentage of IFN+ cells, not the absolute number of transcripts, correlates with viral load in human lung A549 cells, measured by *NP* genomic RNA. **E.** The percentage of IFN+ cells, not the absolute number of transcripts, correlates with viral load in human lung A549 cells, measured by NP protein. **F.** Validation of JAK/STAT signaling inhibitor, Ruxolitnib, in human lung A549 cells. smHCR quantification of *RIG-I* mRNA, which is induced by IFNB treatment for 6 hrs. **G.** Representative smHCR images of influenza-infected human lung A549 cells treated with Ruxolitnib at 6 hpi. **H.** The percentage of IFN+ cells, but not the level of *Ifn* transcripts, positively correlated with viral load in infected A549 cells treated with Ruxolitnib. **I.** IFN production and signaling states among foci at different time points upon infection. **J.** Epithelial-breached foci were larger than epithelial-only foci. **B, C, D, E, H**: r indicates Spearman’s rank correlation coefficient. Asterisk denotes statistical significance. **F, J**: Asterisk denotes statistical significance using one-way ANOVA. **A**, **H**, **J**: Error bars indicate mean +/- SEM.

**Figure S2.**
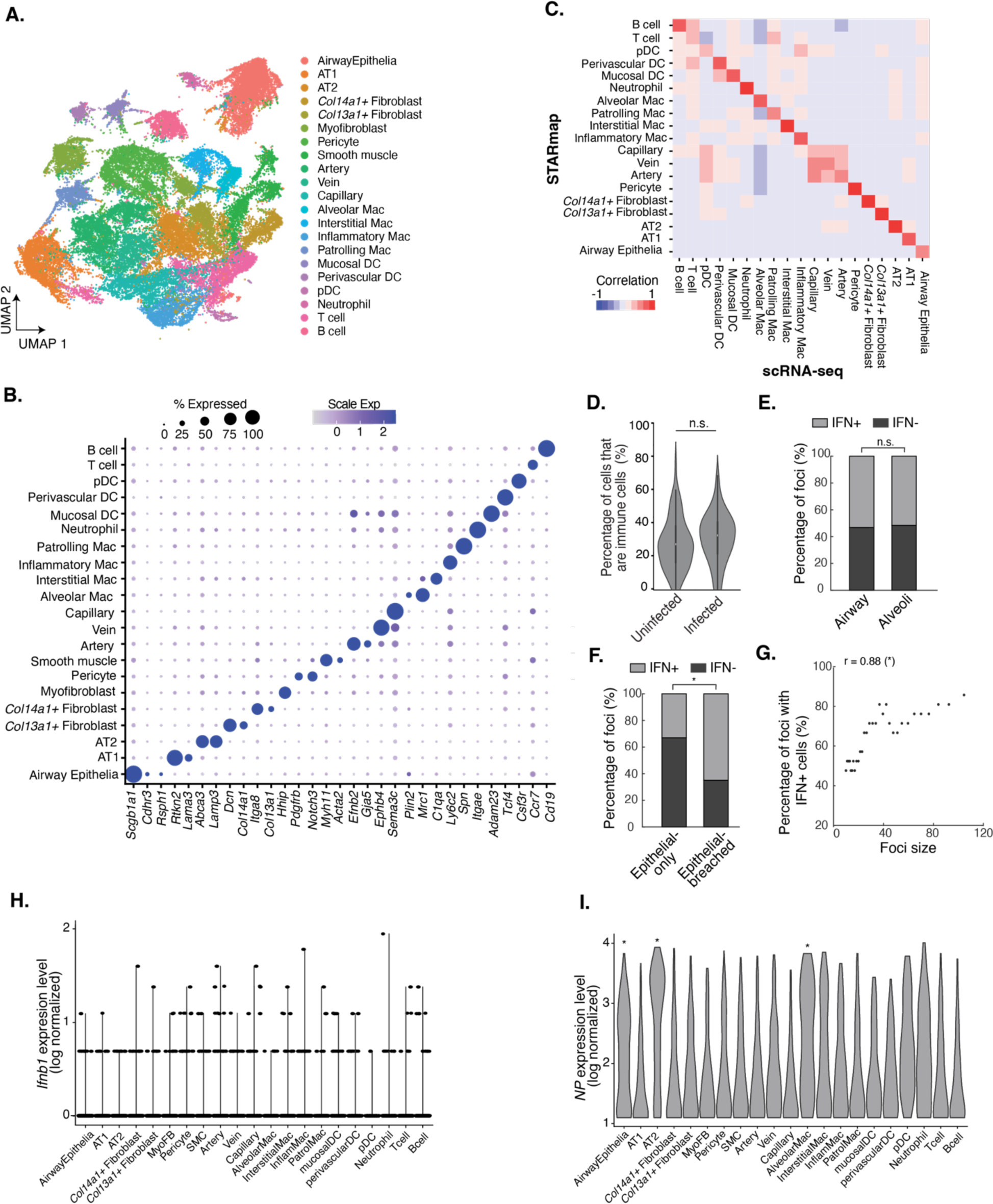
STARmap PLUS reveals differential probability of IFN production among cell types in influenza-infected mouse lung (related to Figure 2). **A.** Uniform manifold approximation and projection (UMAP) of 47,122 cells across 10 lung sections measured via STARmap PLUS. **B.** Dotplot of selected marker gene expression in all identified cell types. **C.** Pairwise Pearson correlation coefficients between the average expression profiles (in z-scores) of individual clusters identified with STARmap PLUS and scRNA-seq ^7^. **D.** No change in the proportion of immune cells found in uninfected neighborhoods vs infected neighborhoods. **E.** The percentage of foci with IFN+ cells was independent of anatomical location. **F.** The percentage of foci with IFN+ cells was higher among epithelia-breached foci than epithelia-only foci. **G.** The percentage of foci with IFN+ cells correlated with focus size. r indicates Spearman’s rank correlation coefficient **H.** Violin plots of IFN level measured by log-normalized *Ifnb1* expression level across all cell types. **I.** Violin plots of viral load measured by log-normalized *NP* expression level across all cell types. **D, E, F, G, I**: Asterisk denotes statistical significance and n.s. denotes no significance using one-way ANOVA (**D**), Spearman’s rank correlation (**G**), Fisher’s exact test (**E**, **F**), Kolmogorov-Smirnov test (**I**).

**Figure S3.**
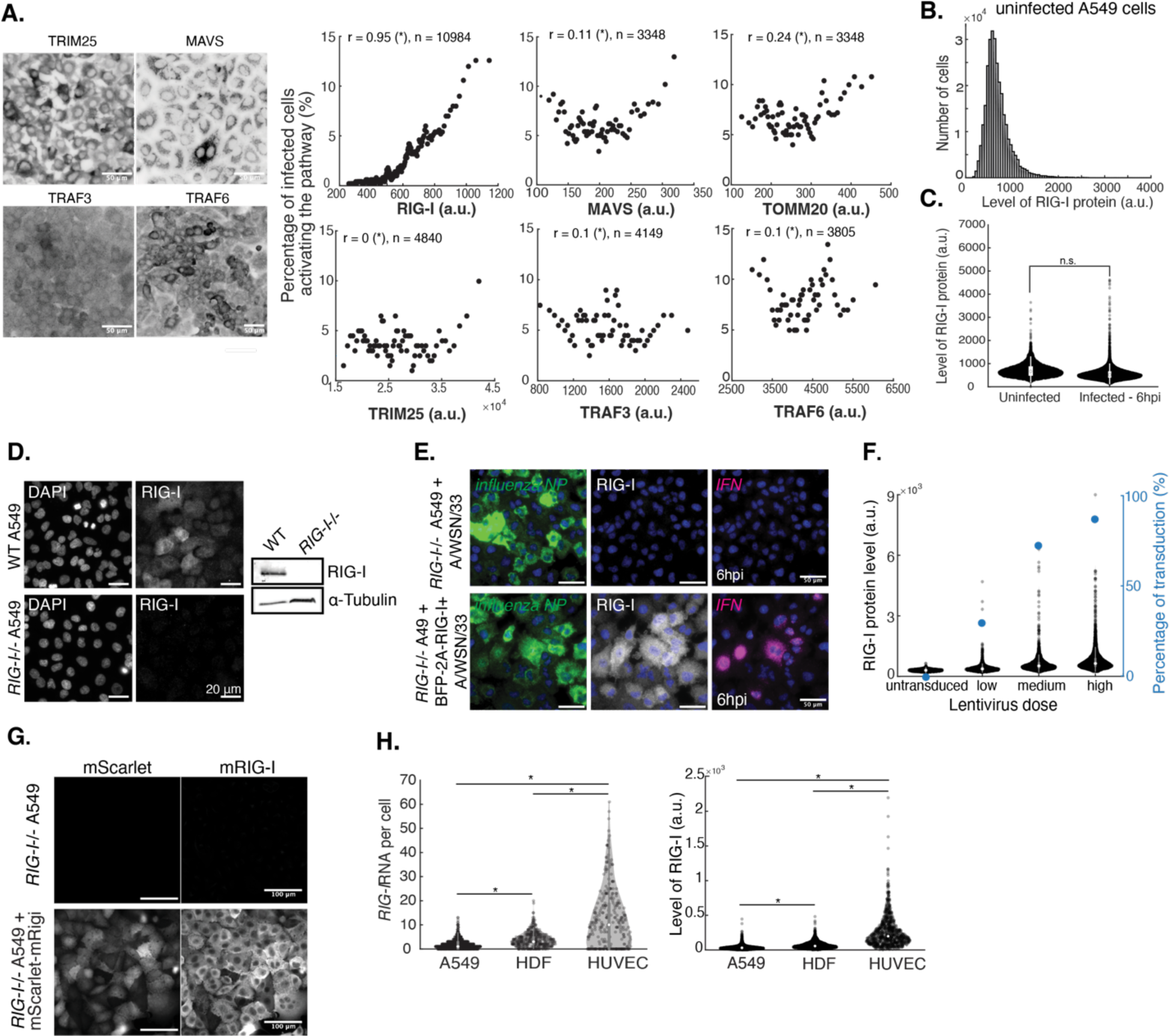
The basal RIG-I level tunes the probability of IFN production upon infection (related to Figure 4). **A.** Immunofluorescence staining on uninfected A549 cells labeled with antibodies against RIG-I pathway components (*left*). The level of RIG-I, but not other pathway components, correlated with the percentage of IFN+ cells (un-normalized protein level) (*right*). r indicates Spearman’s rank correlation coefficient, asterisks denote statistical significance, and n denotes number of cells. **B.** Distribution of RIG-I protein level in uninfected A549 cells. **C.** RIG-I protein level did not change significantly between 0 and 6 hpi. n.s. denotes no significance using one-way ANOVA. **D.** Validation of *RIG-I-/-* A549 cell line. Immunofluorescence staining of RIG-I in wild-type A549 and *RIG-I-/-* A549 (*left*). Western blot of RIG-I and α-tubulin protein in wild-type and *RIG-I-/-* A549 (*right*). **E.** Reconstituting human RIG-I expression in *RIG-I-/-* A549 cells via lentiviral transduction restored stochastic IFN production. **F.** Quantification of RIG-I protein level per cell and the percentage of RIG-I+ cells for each lentiviral dose. **G.** Validation of the antibody against mouse RIG-I. Immunofluorescence staining on *RIG-I-/-* A549 overexpressing mouse RIG-I via lentiviral transduction labeled with antibodies against mouse RIG-I. **H.** Cultured epithelial, fibroblast, and endothelial cell lines have significantly different levels of *RIG-I* mRNA and RIG-I protein, measured by smHCR and immunofluorescence respectively. HDF, human dermal fibroblasts. HUVEC, human umbilical vein endothelial cells. Asterisks denote statistical significance using one-way ANOVA tests. All cells were treated with 10 μm Ruxolitnib in **A, C, E**.

**Figure S4.**
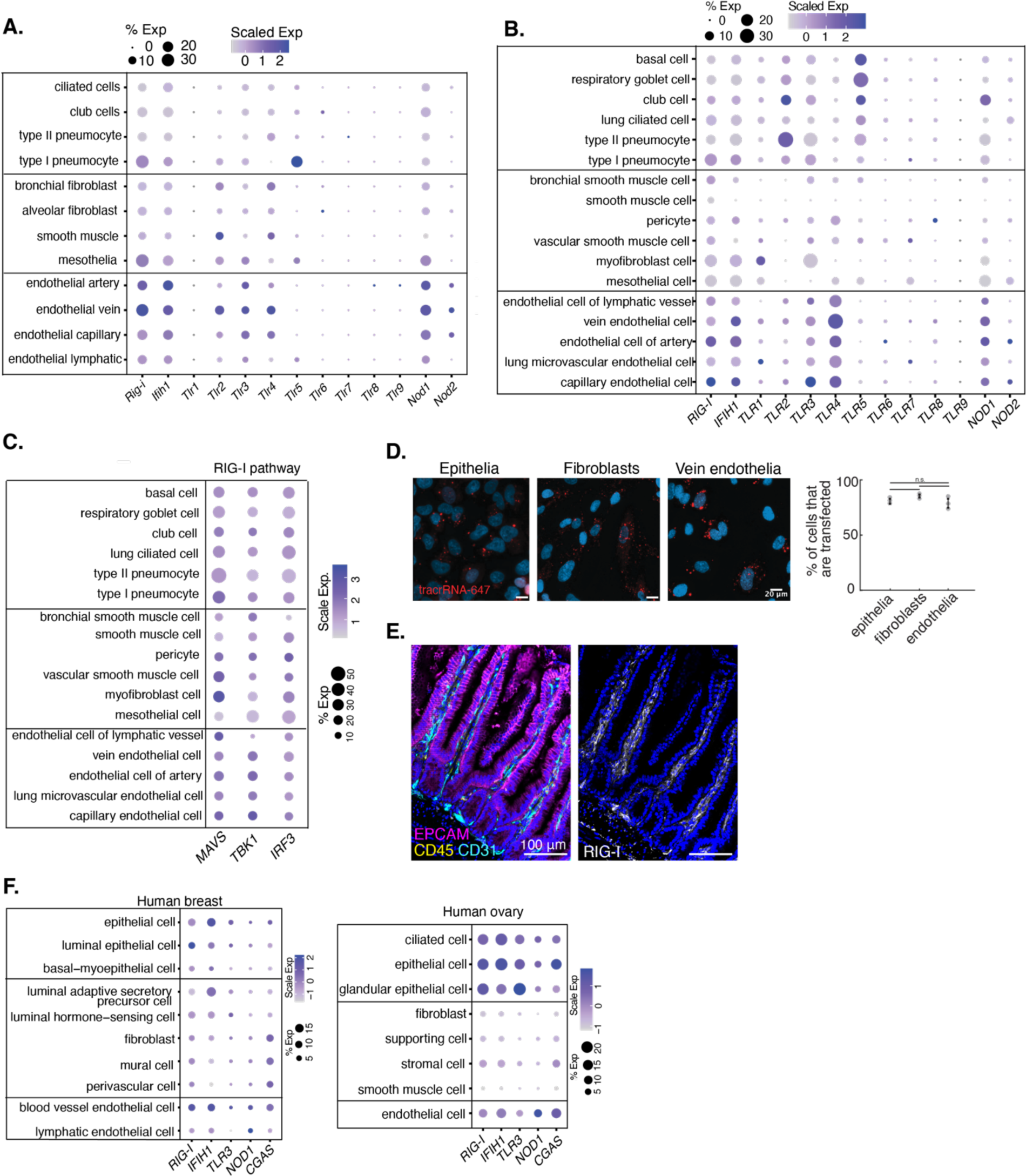
Tiered threat-sensing probability is a general principle of barrier organs (related to Figure 5). **A.** Differential expression of PRRs across cell types in mouse lung. scRNAseq data from Koenitzer et al ^7^. **B.** Differential expression of PRRs across cell types in human lung. scRNAseq data from Tabula Sapiens Consortium ^11^. **C.** Expression of PRR pathway components across cell types in human lung. scRNAseq data from Tabula Sapiens Consortium ^11^. **D.** Validation of transfection efficiency in human primary lung cell types using tracrRNA-Alexa647. Error bars indicate mean +/- SEM. n.s. denotes no significance using one-way ANOVA. **E.** RIG-I protein level was high in immune and endothelial cells and low in epithelial cells in healthy adult intestine. Tissues were labeled with antibodies against EPCAM (epithelia), CD45 (immune), CD31 (endothelia) and and mouse RIG-I. **F.** Expression of PRRs across cell types in non-barrier organs, such as the adult human ovary and human breast ^5,6^.

**Figure S5.**
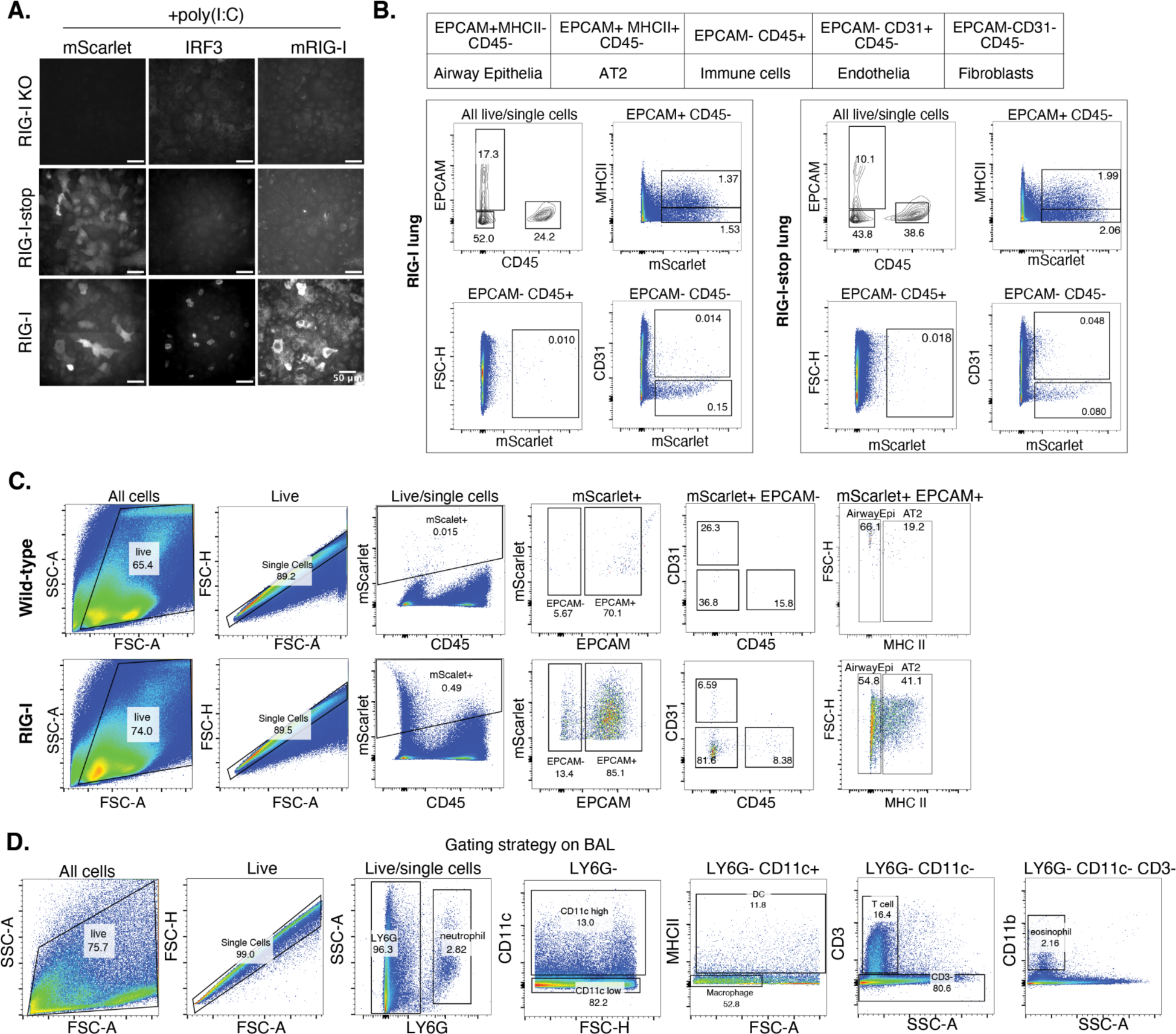
Lentiviral constructs are primarily expressed by lung epithelial cells (related to Figure 6). **A.** Functional validation of RIG-I and RIG-I-stop viruses by transducing *RIG-I-/-* A549 for 24 hrs and subsequently challenging the cells with 250 ng/mL low molecular weight poly(I:C) for 6 hrs. Immunofluorescence staining with antibodies against mouse RIG-I and IRF3. **B.** Gating strategy to identify the cell type composition among mScarlet+ cells 2 weeks post lentiviral transduction. **C.** Gating strategy to identify the transduction efficiency of lentivirus in different cell types 2 weeks post lentiviral transduction. **D.** Gating strategy to identify immune cell types collected in bronchoalveolar lavage.

**Figure S6.**
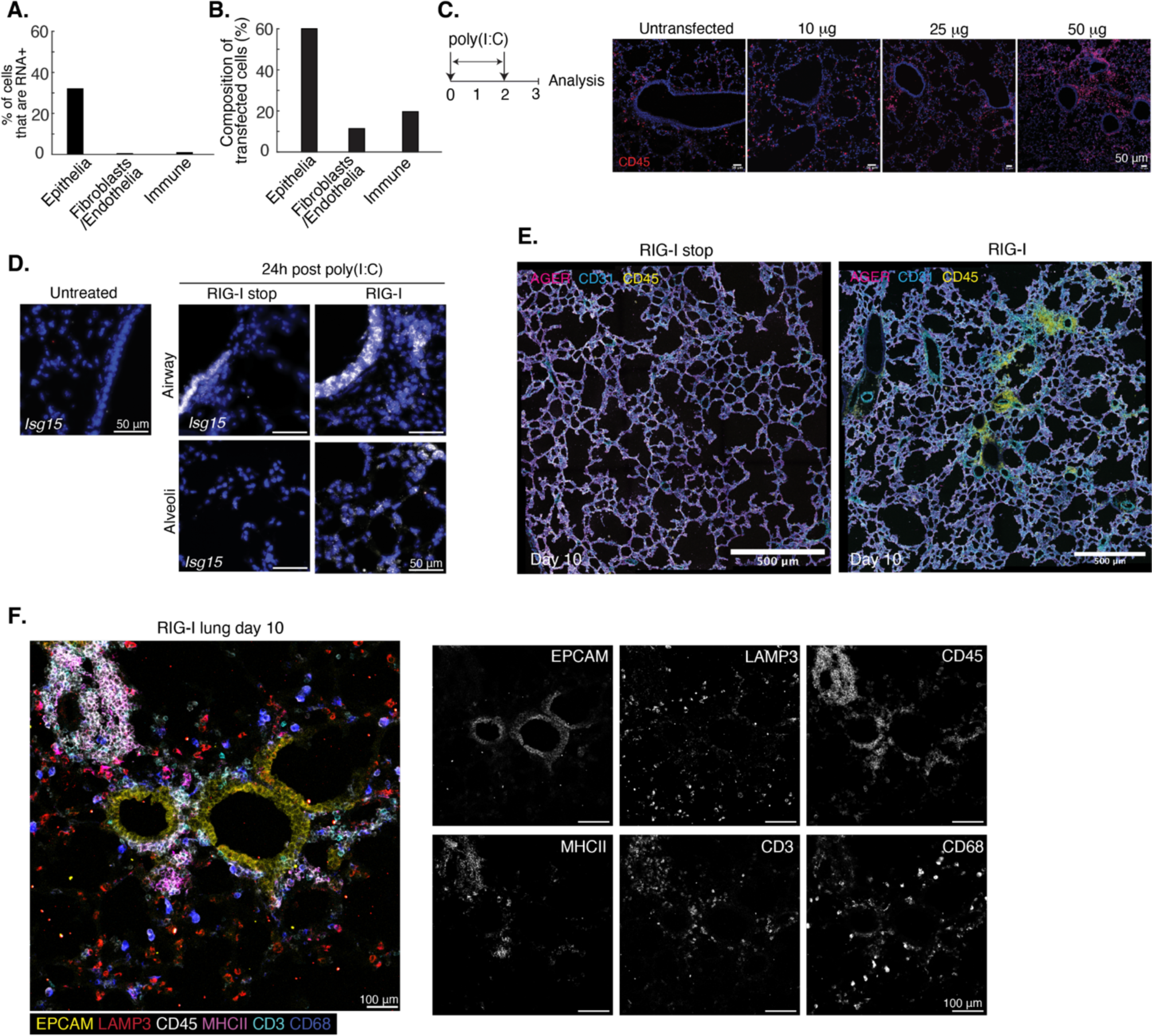
Mice overexpressing RIG-I suffer from more severe tissue damage and prolonged inflammation (related to Figure 6). **A.** Analysis of transfection efficiency by tracrRNA-647 complexed in jetPEI administered intra-nasally. Lung digest was collected after 16 hr and analyzed by flow cytometry. **B.** Composition of transfected cells analyzed by flow cytometry after 16 hr. **C.** Representative images of wild-type mouse lungs upon poly (I:C) challenge at different doses, stained with antibodies against CD45 (immune cells). **D.** Representative smHCR images of RIG-I and RIG-I-stop lung sections labeled with probes against *Isg15* (ISG) at day 1 post poly(I:C). **E.** Mice overexpressing RIG-I had prolonged immune cell infiltration and more severe damage in the alveolar structure at day 10 post poly(I:C). Tissue sections were labeled with antibodies against AGER (AT1), CD31 (endothelia), and CD45 (immune cells) **F.** Composition of immune infiltrates in RIG-I mice at day 10 post poly(I:C). Tissue section was labeled with conjugated antibodies against CD45-Alexa647 (immune cells), MHC-II-PacBlue (antigen-presenting cells), CD3-Alexa488 (T cells), CD68-Alexa594 (macrophages) and unconjugated antibodies against EPCAM (epithelia) and LAMP3 (AT2).

**Supplementary Table 1.** List of genes probed in STARmap PLUS.

| **Cell type markers** | **Ligand/receptors** | **Response genes** |
| --- | --- | --- |
| *Sftpc* | *Ddx58* | *Stat1* |
| *Etv5* | *Ifih1* | *Stat2* |
| *Abca3* | *Dhx58* | *Irf1* |
| *Lamp3* | *Mavs* | *Irf2* |
| *Lama3* | *Tlr3* | *Irf3* |
| *Rtkn2* | *Tlr7* | *Irf7* |
| *Sema3a* | *Ifna2* | *Irf9* |
| *Kcnk2* | *Ifna4* | *Isg15* |
| *Fmo3* | *Ifnb1* | *Isg20* |
| *Scgb1a1* | *Ifnar1* | *Ifit1* |
| *Cldn10* | *Ifnar2* | *Ifitm3* |
| *Cdhr3* | *Ifng* | *Oas1a* |
| *Cfap54* | *Ifngr2* | *Oasl2* |
| *Rsph1* | *Ifngr1* | *Rsad2* |
| *Foxj1* | *Ifnl2* | *Eif2ak2* |
| *Acta2* | *Ifnlr1* | *Gbp2* |
| *Myh11* | *Tnfsf10* | *Gbp3* |
| *Lmod1* | *Tnfrsf10b* | *Bst2* |
| *Dcn* | *Tradd* | *Cd74* |
| *Col14a1* | *Tnfrsf21* | *B2m* |
| *Prrx1* | *Fas* | *Trim30a* |
| *Mmp3* | *Fasl* | *Trim25* |
| *Col13a1* | *Tnf* | *Irgm1* |
| *Itga8* | *Tnfrsf1a* | *Cd274* |
| *Npnt* | *Tnfrsf1b* | *Nlrp3* |
| *Hhip* | *Cxcl1* | *Usp18* |
| *Aspn* | *Cxcl2* | *Socs1* |
| *Adamts17* | *Cxcl3* | *Gbp4* |
| *Pdgfrb* | *Cxcl5* | *Gbp7* |
| *Notch3* | *Cxcr1* | *Pias1* |
| *Postn* | *Cxcr2* | *Samhd1* |
| *Efnb2* | *Cxcl9* | *Traf2* |
| *Sox17* | *Cxcl10* | *Traf1* |
| *Gja5* | *Cxcl11* | *Nfkb1* |
| *Ephb4* | *Cxcr3* | *Rela* |
| *Vegfc* | *Ccl2* | *Birc2* |
| *Nrp2* | *Ccl7* | *Birc3* |
| *Prox1* | *Ccl12* | *Nfkbia* |
| *Ear2* | *Ccr2* | *Tnfaip3* |
| *Plin2* | *Ccl3* | *Zfp36* |
| *Plet1* | *Ccl4* | *Dusp8* |
| *Siglecf* | *Ccl5* | *Prdm1* |
| *C1qa* | *Ccl11* | *Timp1* |
| *C1qb* | *Ccr5* | *Stat3* |
| *Mrc1* | *Ccr3* | *Ppara* |
| *Ly6c2* | *Csf2* | *Socs3* |
| *Ccr2* | *Csf2rb* | *Pias3* |
| *F13a1* | *Il6* | *Ptpn6* |
| *Spn* | *Il6ra* | *Myd88* |
| *Dusp16* | *Il6st* | *Traf6* |
| *Nr4a1* | *Il1a* | *Il1rl1* |
| *Etv3* | *Il1b* | *Smad7* |
| *Tnip3* | *Il1r1* | *Psmb9* |
| *Adam23* | *Il17a* | *Atf3* |
| *Itgae* | *Il17ra* | *Nfil3* |
| *Arsb* | *Il18* | *Hmox1* |
| *Plbd1* | *Il18r1* | *Blvra* |
| *Siglech* | *Il18bp* | *Blvrb* |
| *Grm8* | *Il4* | *Smad3* |
| *Lrp8* | *Il4ra* | *Cdkn1a* |
| *Tcf4* | *Il2* | *Col1a2* |
| *Ly6g* | *Sema3c* | *Thbs1* |
| *Csf3r* | *Plxnd1* | *Itga2* |
| *Nrg1* | *Vegfa* | *Casp8* |
| *Tcf7* | *Kdr* | *Casp9* |
| *Ccr7* | *Fgf2* | *Casp3* |
| *Gzmb* | *Fgfr1* | *Casp7* |
| *Tbx21* | *Tgfb1* | *Casp1* |
| *Gata3* | *Tgfbr1* | *Apaf1* |
| *Rorc* | *Tgfbr2* | *Runx3* |
| *Foxp3* | *Il10* | *Bcl2* |
| *Ctla4* | *Il10ra* | *Bcl2l1* |
| *Il2ra* | *Cd200* | *Mcl1* |
| *Pax5* | *Cd200r1* | *Cflar* |
| *Bank1* | *Itga5* | *Ubc* |
| *Cd19* | *Itgb6* |  |
| *Influenza NP* | *Vcam1* |  |
| *Influenza NS1* | *Itgb2* |  |
|  | *Itgam* |  |
|  | *Icam1* |  |
|  | *Icam2* |  |
|  | *Sele* |  |
|  | *Adam17* |  |
|  | *Adam10* |  |

**Supplementary Table 2.**
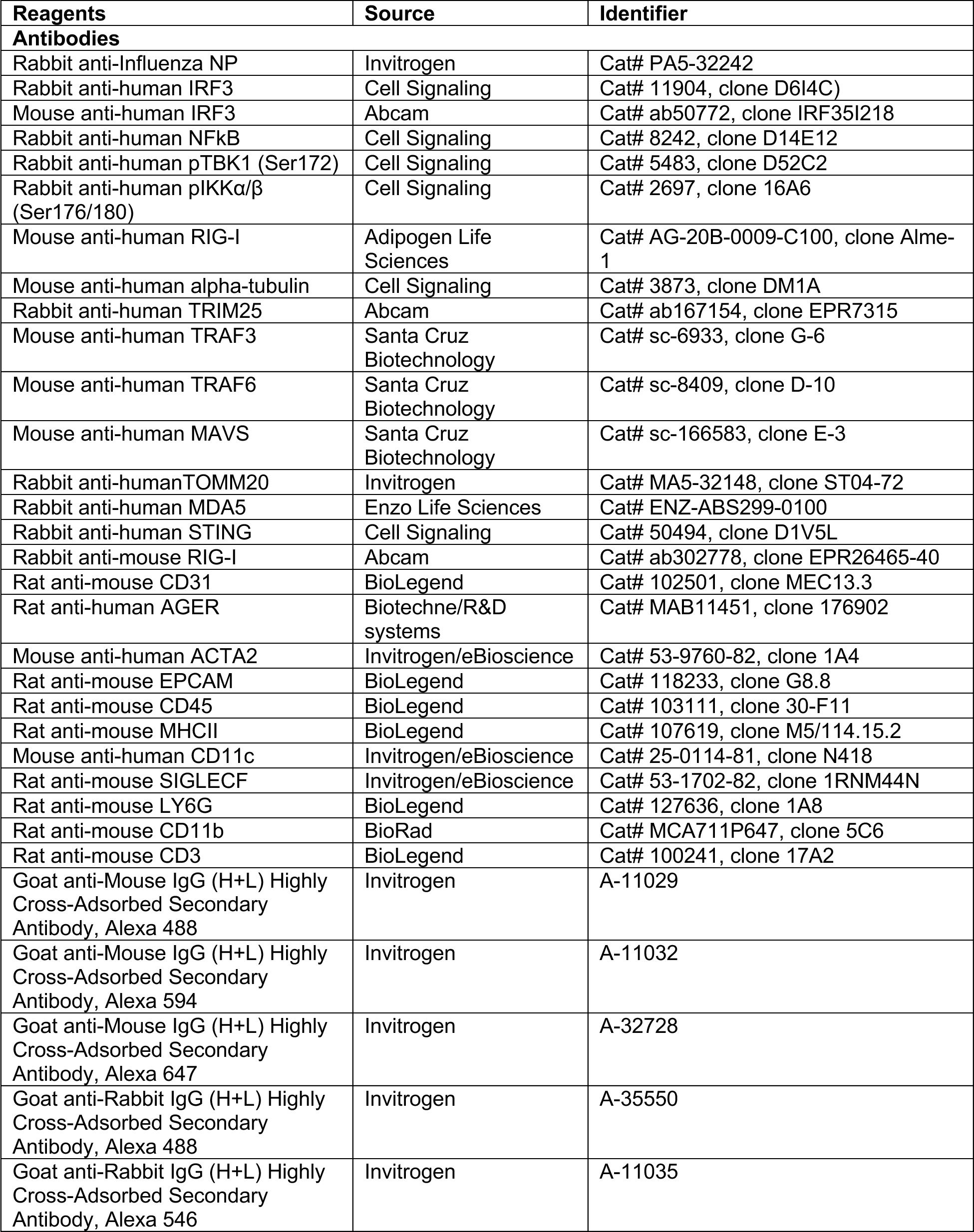
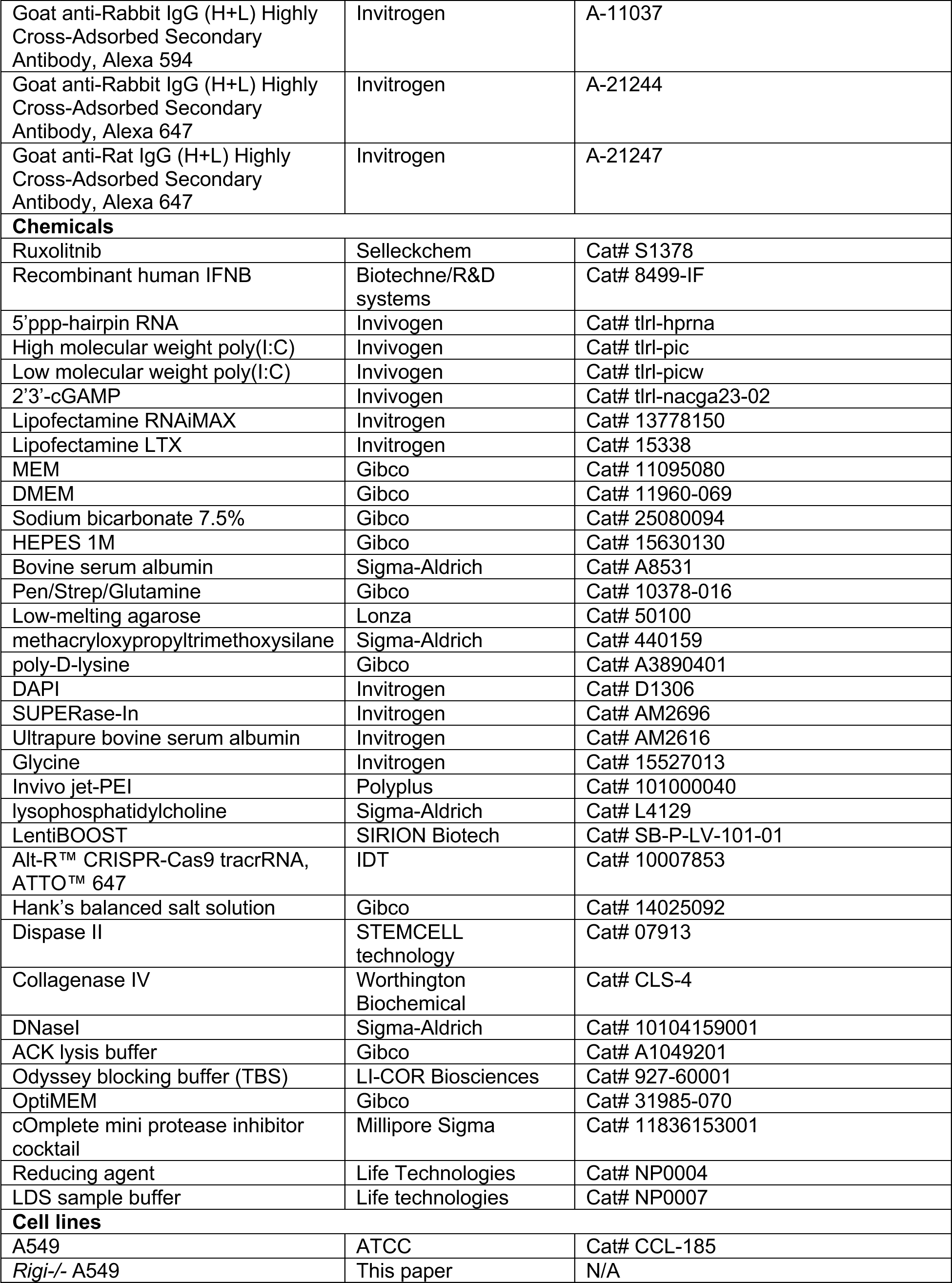
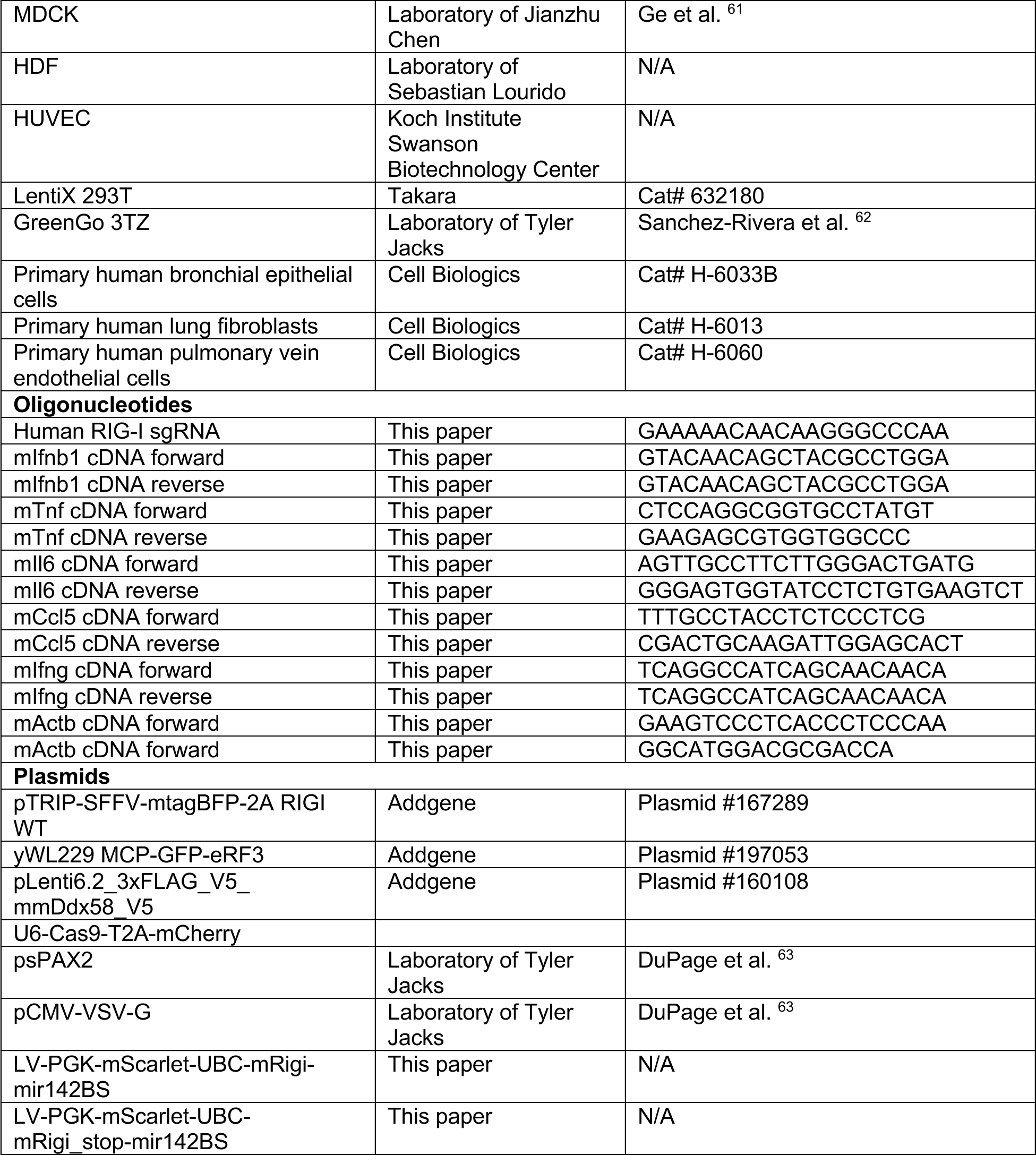
List of key reagents.

